# Task-specific representational spaces for memory formation in the medial temporal lobe

**DOI:** 10.1101/2025.05.23.655754

**Authors:** Elias M.B. Rau, Rebekka Heinen, Nora A. Herweg, Eva Z. Patai, Malte Kobelt, Khazar Ahmadi, Nikolai Axmacher

## Abstract

Adaptive behavior requires stimuli to be processed in a way that meets the requirements of the current task. This is especially relevant for natural concepts that comprise a multitude of representational formats. The medial temporal lobe (MTL) has been proposed to support such flexibility by organizing stimuli within relational representational spaces serving as cognitive maps. However, while various studies investigated the formation of novel cognitive maps based on abstract features, it remains unclear how our vast existing conceptual knowledge is recruited into task-relevant representational spaces and whether these spaces influence the formation of novel episodic memories. Here we show that neural representations of natural concepts in the MTL and neocortex are embedded in low-dimensional representational spaces of task-relevant features that flexibly reorganize when task demands change. Crucially, hippocampal pattern similarity reflects task-relevant, but not task-irrelevant conceptual feature dimensions of the same set of natural concepts, indicating context-dependent reconfiguration of representational spaces. Euclidean distances within and across these spaces predicted subsequent memory confidence for individual items, suggesting a putative metric of task-dependent neural distinctiveness for the formation of memory traces. Finally, we dissociate task- and stimulus-dependent formats in neocortex, demonstrating complementary representations of natural stimuli that differentially contribute to memory formation. Our findings suggest that relational coding of task-relevant features in the MTL may serve as an index of representational distinctiveness that contributes to the formation of novel episodic memories and integrate MTL accounts based on spatial coding and episodic memory.

## Introduction

Natural stimuli are processed in a multitude of representational formats in the brain. These formats may reflect their physical appearance or semantic category but also context-dependent formats dictated by behavioral demands (Hebart et al. 2020; Heinen et al. 2023, 2025a). Consider a farmer and one of their cows: the same cow may be judged by strikingly different features depending on the situation, for example when planning a seasonal grazing schedule as compared to evaluating its value on the market. Although the sensory input and the recognized object are identical, the behaviorally relevant information is abstract, memorized, and dictated by context rather by properties of the specific stimulus. Importantly, such task-dependent differences in the relative weights of concept features have implications for a concepts’ relationship towards related concepts in feature-specific representational space (Tversky 1977), a property that can be measured via behavioral distance judgements and neural pattern similarity. Importantly, such context-dependent flexibility is also crucial for episodic memory, which requires the separation of different memories with overlapping elements (Yassa and Stark 2011).

Here, we test the hypothesis that the behavioral flexibility in judging concepts differently between contexts is enabled by neural representations in the MTL via embeddings of concepts in task-specific representational feature spaces. Specifically, akin to MTL representations of cognitive maps (Epstein et al. 2017), these embeddings should be organized by behaviorally relevant feature dimensions, express relationships between embedded concepts, and reorganize flexibly when task demands change – properties of neural representations that can be studied using representational similarity analysis (Kriegeskorte 2008). We therefore ask 1) how the brain encodes such task-specific representational formats, 2) how properties of concept relationships in task-relevant representational spaces are associated with the formation of novel episodic memories, and 3) how task- and stimulus-dependent representational formats contribute to successful encoding.

Perceptual formats can be studied using convolutional deep neural networks (DNNs), which capture increasingly complex visual features of neural representations along the processing hierarchy of the ventral visual stream (VVS) (Cichy et al. 2016; Güclü, and van Gerven 2015; Kuzovkin et al. 2018) and have more recently been applied to mnemonic representations (Davis et al. 2021; Heinen et al. 2025b; Rau et al. 2025). Conceptual formats, as e.g. captured in linguistic descriptions, have been related to neural representations in both VVS and anterior temporal cortices (Devereux, Clarke, and Tyler 2018; Huth et al. 2012, 2016), and converging evidence supports the notion that such perceptual and conceptual formats can be supported by multiple parallel representational spaces in the brain (Deuker et al. 2016; Zheng et al. 2024). Beyond stimulus-dependent properties, representing context-dependent information requires top-down modulation of stimulus-driven codes, as suggested by evidence that task context biases sensory representations towards high-level, behaviorally relevant information (Shin, Doron, and Larkum 2021; Nigam and Schwiedrzik 2024).

A growing body of evidence suggests that the hippocampus (HC) and medial temporal lobe (MTL) selectively represent information relevant to the current behavioral context. For example, top-down attention modulates neural representations of the same stimulus material (Aly and Turk-Browne 2016b, 2016a; Nobre and Stokes 2019) and HC representational geometry changes as a function of learning and updated task goals (Mack, Love, and Preston 2016; Theves, Fernandez, and Doeller 2019; Theves, Fernández, and Doeller 2020). Context-dependent representations in the MTL are further supported by studies on mixed selectivity — flexible changes in neuronal responsiveness depending on task demands (Fusi, Miller, and Rigotti 2016; Han et al. 2023) — and by HC place cell remapping in rodents, whereby cells change their responses between contexts, enabling orthogonalized, context-dependent memory representations (Fyhn et al. 2007; Colgin, Moser, and Moser 2008; Kubie, Levy, and Fenton 2020). A compelling framework that describes such flexible, relational coding are cognitive maps, first introduced by Tolman (1948), who formalized the idea of a common representational space organized by metric distances, that allows to infer previously unexperienced trajectories. Indeed, neurophysiological evidence for cognitive maps comes from the discovery of grid cells in entorhinal cortex (EC) (Fyhn et al. 2008; Jacobs et al. 2013), and HC place cells (O’Keefe 1976). Further, HC neurons in humans have been found to code for distinct concepts (Quiroga et al. 2005, 2008; Rey et al. 2015) as well as their conjunctions with episodic context (Kolibius et al. 2023), suggesting a more general role in relational coding.

Accordingly, cognitive maps and their neural basis have been proposed to generalize beyond spatial navigation to the organization of knowledge in memory (Behrens et al. 2018; Bellmund et al. 2018; Eichenbaum 2017), suggesting that low-dimensional representational spaces organized by task-relevant features may be a domain-general principle of MTL coding that is recruited during spatial navigation but also for the dynamic scaffolding of individual items and concept knowledge (Whittington et al. 2020). Such task-specific relational structure can be tracked in macroscopic brain signals via RSA as demonstrated in prior fMRI studies, which showed that hippocampal representational distances correspond not only to spatial distances (Deuker et al. 2016; Hassabis et al. 2009) but also to perceptual (Viganò et al. 2021), associative (Constantinescu, O’Reilly, and Behrens 2016; Theves et al. 2019, 2020), and abstract features (Garvert et al. 2023; Garvert, Dolan, and Behrens 2017; Tavares et al. 2015; Teoh et al. 2026).

However, most of these studies investigated the formation of novel cognitive maps via explicit associative learning of artificially constructed stimulus-feature pairings, leaving open whether such map-like representational spaces flexibly organize naturally acquired concept knowledge. Moreover, apart from the learning of the relational structure itself, the use of repeated stimulus exposure during acquisition precluded any link between relational encoding and the formats of novel episodic memories. The present study addresses these gaps. First, we capitalize on participants’ existing knowledge about animal concepts by obtaining pairwise distance ratings along natural feature dimensions that do not require explicit, experimentally controlled learning. Second, the same concepts are rated in two distinct behavioral contexts, enabling direct within-concept comparison — a macroscopic analog of context-dependent coding — to test whether HC representational geometry is flexibly reorganized by task demands. Third, we examine whether relational metrics within and across task-specific representational spaces are associated with subsequent memory strength for individual exemplars of a concept, a putative index to reconcile between spatial and episodic accounts of the medial temporal lobe in human memory processes.

## Results

### Task-dependent encoding of natural concepts

While undergoing fMRI, participants engaged in a conceptual distance rating task that involved the viewing of natural animal images and subsequent ratings based on conceptual feature dimensions (Heinen et al. 2025b). In each trial, participants viewed two trial-unique exemplars of animals and judged their relationship according to two feature dimensions (e.g. *Size* & *Climate*, or *Proximity* & *Activity*; see Fig. 1A; fig. S1) by rating the distance of the second animal relative to the first animal on a 10-point scale. Both rated features were instructed at the beginning of each block, but the pairing of features (e.g. *Size* & *Climate*) was constant within a participant. Importantly, each animal concept appeared equally often in each block (with different exemplars in different blocks), allowing direct comparisons between its relationship to other animals in two task contexts. The resulting pairwise distance ratings were then used to reconstruct task-specific representational spaces of animal relationships, and these spaces were compared to neural activity patterns during image presentations. This allowed us to test whether the representational geometry of concepts was behaviorally relevant and whether it depended on task demands (for details see Methods).

**Figure 1.**
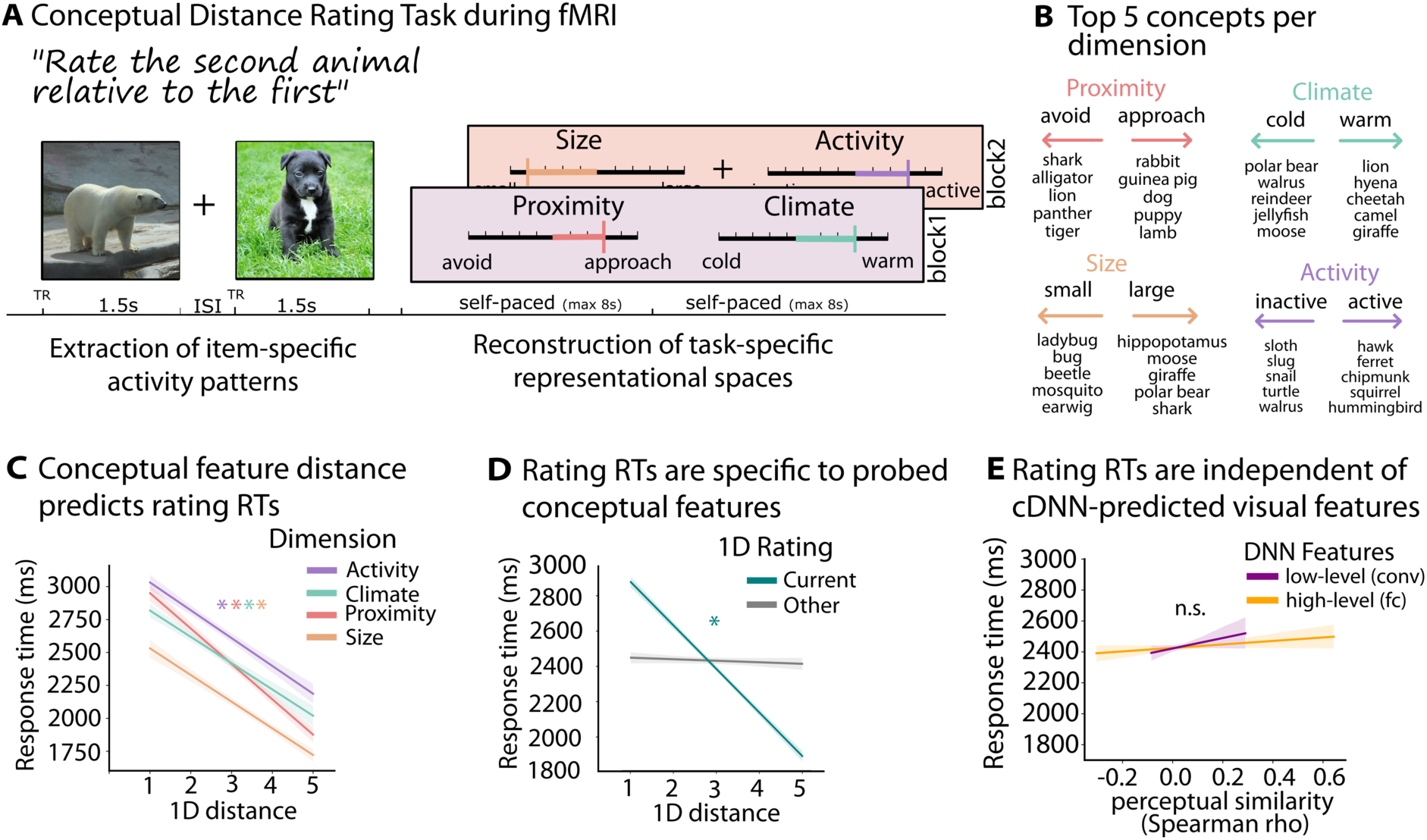
Conceptual distance ratings are specific to natural feature dimensions. A) Exemplary trial of the conceptual distance rating task performed during fMRI scanning. Image onset events were used to extract image-specific neural activity patterns. Rating events were used for reconstruction of representational spaces. B) Top 5 animal concepts for each conceptual feature dimension and pole based on grouped responses across participants (*Proximity*: avoid – approach; *Climate*: cold – warm; *Size*: small – large; *Activity*: inactive – active; for details see Methods). C) Magnitude of rated distance (1D) significantly predicts RTs in each conceptual feature dimension. D) RTs are predicted by distance rating of currently probed feature dimension (teal), but not by the other feature dimension rated in the same trial (grey). E) Perceptual similarity between image exemplars in a trial (low-level perceptual: convolutional DNN layers; high-level perceptual: fully-connected DNN layers) are not related to rating RTs. * indicates beta-coefficient with *P* < 0.05 after model comparison.

We first verified that distance ratings reflected task demands and that the rated distances meaningfully organized animals along each conceptual feature dimension (Fig. 1B). Using linear mixed models (LMMs), we found that response times (RTs) were shorter when animals differed more on the rated feature dimension (all *Z* < −13.253, all *χ*^2^ > 171.60, all *P* < 0.001; Fig. 1C). Specifically, lower RTs were predicted by the currently probed conceptual feature dimension (current feature: *Z* = −35.628, *χ*^2^ = 1217.44, *P* < 0.001), but not by the other feature dimension probed in the same trial (other feature dimension: *Z* = 23.107, *χ*^2^ = 10.89, *P* = 0.001), suggesting specificity of responses to different conceptual features within a trial (Fig. 1D). Further, RTs were independent of the perceptual similarity between both image exemplars, as predicted by a convolutional DNN (“AlexNet”; Krizhevsky et al. 2017) that captures image similarity estimates based on low-level (convolutional layers) and high-level (fully-connected layers) visual features (convolutional layers: *Z* = 1.088, *χ*^2^ = 1.18, *P* = 0.277; fully-connected layers: *Z* = 0.967, *χ*^2^ = 0.94, *P* = 0.333; Fig. 1E; for details see Methods). These results confirm that distance ratings and response times depended on the task-relevant conceptual properties but not on general stimulus-related properties of individual exemplar images.

### Reconstruction of task-specific representational spaces

Throughout the experiment, we explicitly sampled distance ratings for ∼10 % of all possible concept pairs. However, to allow inferences to unseen concept pairs, a requirement to investigate relational embeddings in representational spaces, we reconstructed the relative positions for each concept and feature dimension separately using least-squares solutions (Fig. 2A; fig S1; for details see Methods). Following this procedure, we reconstructed two participant-specific two-dimensional representational spaces, one space for each pairing of conceptual feature dimensions (Fig. 2B). These reconstructions have several advantages: They provide a reconstructed Euclidean distance metric between all concepts pairs (explicitly rated and inferred), allow contrasts between task-relevant vs. task-irrelevant distances (i.e. distances between animals based on the currently probed features vs. distances between the same animals based on the features probed in a different block) and across-space comparisons in individual concept positions.

**Figure 2.**
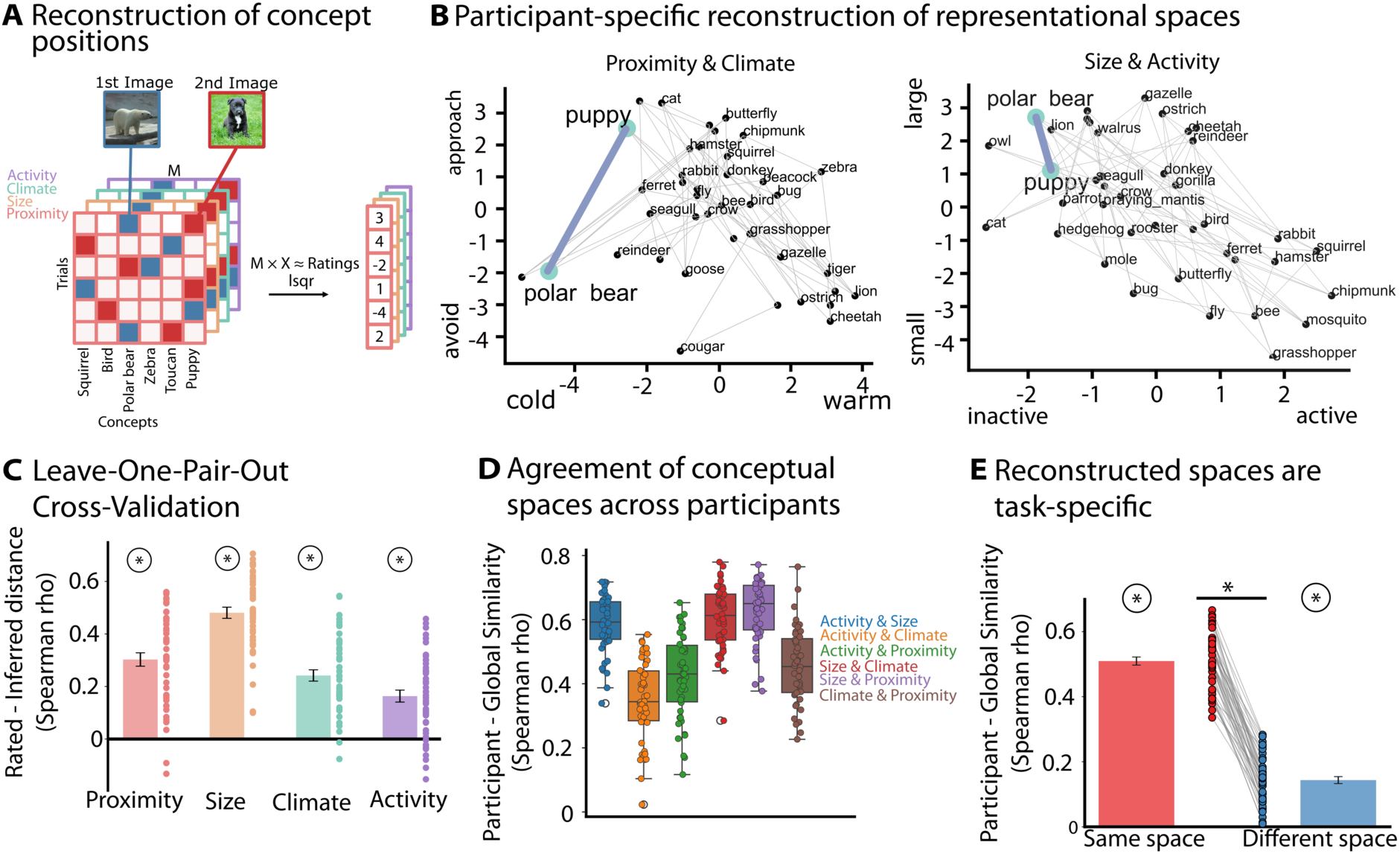
Reconstruction of task-specific representational spaces. A) Reconstructing distances between all concept pairs (rated and inferred), separately for each conceptual feature dimension, using least-squares to approximate behavioral ratings (for details see Methods). B) Exemplary participant-specific reconstruction of 2D representational spaces. Positions for each concept and axis were estimated based on the procedure depicted in A) and combined into the two 2D spaces rated by the participant. Grey lines between concepts indicate pairs of concepts that were explicitly rated in the conceptual distance rating task. Bold lines indicate exemplary distance for the concept pair shown in A), i.e. puppy and polar bear. C) Results of leave-one-pair-out cross-validation between rated and inferred distances. Feature-specific distributions suggest that empirically obtained distances can be meaningfully predicted using left-out data. D) Spearman correlations between participant-specific and global representational spaces separately for each of the six possible spaces. E) Spearman correlations between participant-specific and global representational spaces for spaces with shared feature dimensions (same space) and different feature dimensions (different space). Circled * indicates *P* < 0.05 for one-sample *T* test and lined * indicates *P* < 0.05 for paired *T* test.

We tested the validity of reconstructed spaces by ensuring that the obtained least-squares solutions predicted behavioral ratings significantly better than chance (permutation test for each dimension, all *P* < 0.001; fig. S1; for details see Methods). Second, we conducted a leave-one-pair-out cross-validation to demonstrate that inferred distances meaningfully reflect empirically obtained behavioral distance ratings. Comparisons between inferred and explicitly sampled distances yielded significant positive correlations across participants and for each conceptual feature dimension (Fig. 2C; all *T*_45_ > 7.185, all *P* < 0.001, mean *ρ* = 0.297 ± 0.153), suggesting that distances from reconstructed representational spaces allow generalizations to unseen concept pairs.

As a prerequisite for investigating task-specificity of representational spaces, we first established that animals probed in different spaces yielded a distinct and reliable representational embedding of concept relationships. To do so, we estimated the similarity between participant-specific spaces and space reconstructions based on the ratings of the remaining sample (global; leave out one participant; for details see Methods; for global space reconstructions see fig. S2). Indeed, Participant-global similarity was significantly larger than zero for all spaces (all *T*_45_ > 18.462, *P* < 0.001; mean across spaces *ρ* = 0.507±0.076; Fig. 2D), suggesting a shared relational structure across participants. Next, we computed the similarity of each participant-specific representational space with its matching global space (“task-relevant space”) and the non-matching global space (“task-irrelevant space”; i.e. the space that the participant rated in the other task block). Similarities were significantly larger for task-relevant as compared to task-irrelevant spaces (same space: *ρ* = 0.510±0.085; different space: *ρ* = 0.144±0.073; *T*_45_ = 29.209, *P* < 0.001; Fig. 2E), confirming that concept embeddings were specific to the currently probed feature dimensions.

### Task-specific representational spaces in the medial temporal lobe

Next, we sought to investigate the neural basis for task-dependent representational spaces in the brain, and tested the hypothesis that the MTL, and in particular the hippocampus, dynamically represent concepts according to current task demands. Specifically, in a ROI-based RSA, we tested whether the pairwise relationships between concepts derived from task-relevant representational spaces corresponded to multi-voxel pattern similarity in the brain (for details see fig. S3). We extracted image-specific beta maps, controlled pairwise similarity estimates for mean BOLD magnitude and cDNN-predicted visual features, and averaged across repetitions of the same concept, separately for each space (for details see Methods). We restricted the RSA to those concept pairs that were never directly rated in the experiment (≈90% of concept pairs inferred) but whose relationships were inferred from behaviorally reconstructed spaces. Next, we correlated neural concept embeddings with relationships predicted by behaviorally reconstructed spaces, separately for matching and non-matching space pairs (Fig. 3A). We found that the neural similarity in HC (*T*_46_ = 2.455, *P* = 0.018) and PRC (*T*_46_ = 2.654, *P* = 0.011), but not in PHC (*T*_46_ = 1.823, *P* = 0.075) or EC (*T*_46_ = 0.261, *P* = 0.795) correlated with distances from task-relevant representational spaces (Fig. 3B), suggesting that concept relationships that support current task demands are reflected in MTL neural pattern similarity and importantly, that these neural concept representations generalize beyond explicitly probed comparisons, suggesting embeddings in generalized conceptual spaces. We also observed significant negative correlations between neural pattern similarities and concept relationships predicted by the task-irrelevant representational space in PRC (Fig. 3C; *T*_46_ = −3.582, *P* = 0.001) and PHC (*T*_46_ = −3.379, *P* = 0.002) but not HC (*T*_46_ = −0.252, *P* = 0.802) or EC (*T*_46_ = 0.102, *P* = 0.919). Most importantly, we found that neural representations were task-specific, i.e. significantly more aligned with distances from task-relevant as compared to task-irrelevant representational spaces in HC (*T*_46_ = 2.320, *P* = 0.025), PRC (*T*_46_ 4.512, *P* < 0.001), and PHC (*T*_46_ = 3.646, *P* = 0.001), but not in EC (*T*_46_ = 0.139, *P* = 0.890), suggesting distinct and task-specific representations of the same natural concepts in the MTL. These findings were confirmed in a searchlight-based RSA analysis within regions of the medial temporal lobe, which yielded clusters of significant voxels located in perirhinal cortex and HC that exhibit substantial correlations between neural and reconstructed concept relationships, both for task-relevant spaces but importantly also for task-specific contrasts (i.e. relevant vs. irrelevant spaces; Fig. 3C; Table S1). Task-dependent effects separately for types of rated spaces (e.g. *Activity* & *Size*, *Proximity* & *Climate*) and MTL ROIs can be found in fig. S4.

**Figure 3.**
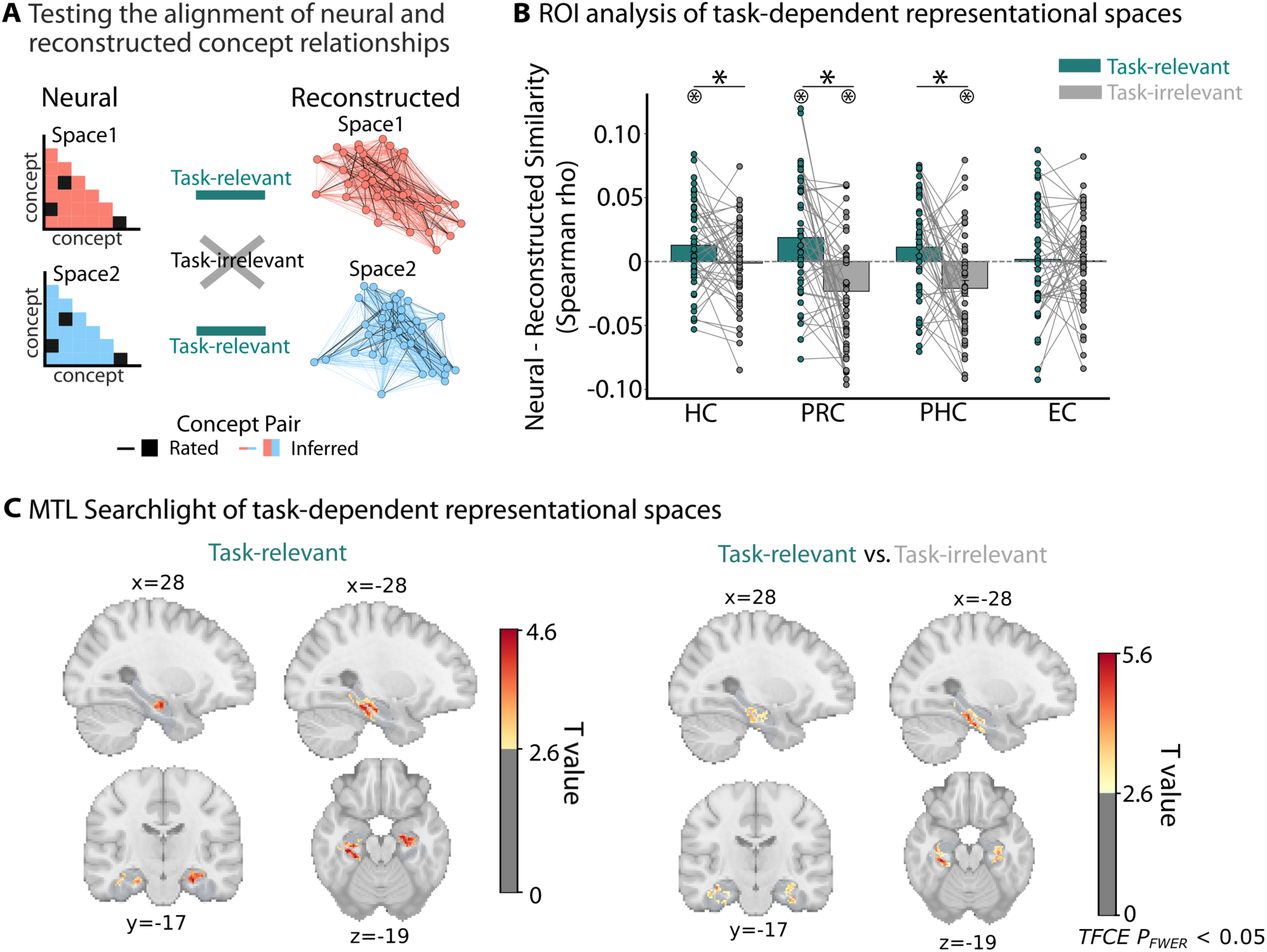
Task-specific representational spaces in the medial temporal lobe. A) Analysis strategy to test the alignment of neural and reconstructed animal concept relationships. We correlated neural similarities between concepts with reconstructed representational geometries of task-relevant and task-irrelevant spaces based on distances of inferred concept pairs (task-relevant: neural_space1_-behavioral_space1_ and neural_space2_-behavioral_space2_; task-irrelevant: neural_space1_-behavioral_space2_ and neural_space2_-behavioral_space1_) and averaged each condition across spaces. Only inferred concept relationships were included (i.e. those not explicitly rated by a participant). B) ROI-based results of representational alignment between neural and reconstructed relationships for task-relevant (teal) and task-irrelevant (grey) spaces across regions of the MTL. C) Same analysis as described in A) but conducted using an MTL searchlight, for task-relevant (top) and task-specific (bottom; i.e. task-relevant vs. task-irrelevant) contrasts. All depicted clusters survive TFCE-based correction for multiple comparisons at the family-wise error rate. Circled * indicates *P* < 0.05 for one-sample *T* test and line * indicates P < 0.05 for paired *T* test.

### Flexible reorganization of task-specific representations in the hippocampus

Our findings so far show that patterns of activity in the MTL correspond to task-specific representational spaces. However, it is unclear whether flexible changes of these neural representations match the reorganization of concept relationships afforded by changes in task demands. To address this question, we correlated similarities between task-dependent representational spaces (space_1_-space_2_ similarity) in MTL subregions with the corresponding similarities of the reconstructed representational spaces. Indeed, across participants, we found a significant positive correlation between the similarity of reconstructed and neural spaces in the HC (*ρ* = 0.405, *P* = 0.005; Fig. 4A; fig. S5), but not in any other MTL region (all *ρ* < 0.212, all *P* > 0.15), suggesting 1) that spaces with similar relational embeddings of concepts are also represented more similarly in the brain and 2) that the HC specifically contributes to this flexible reorganization of representational spaces according to task demands.

**Figure 4.**
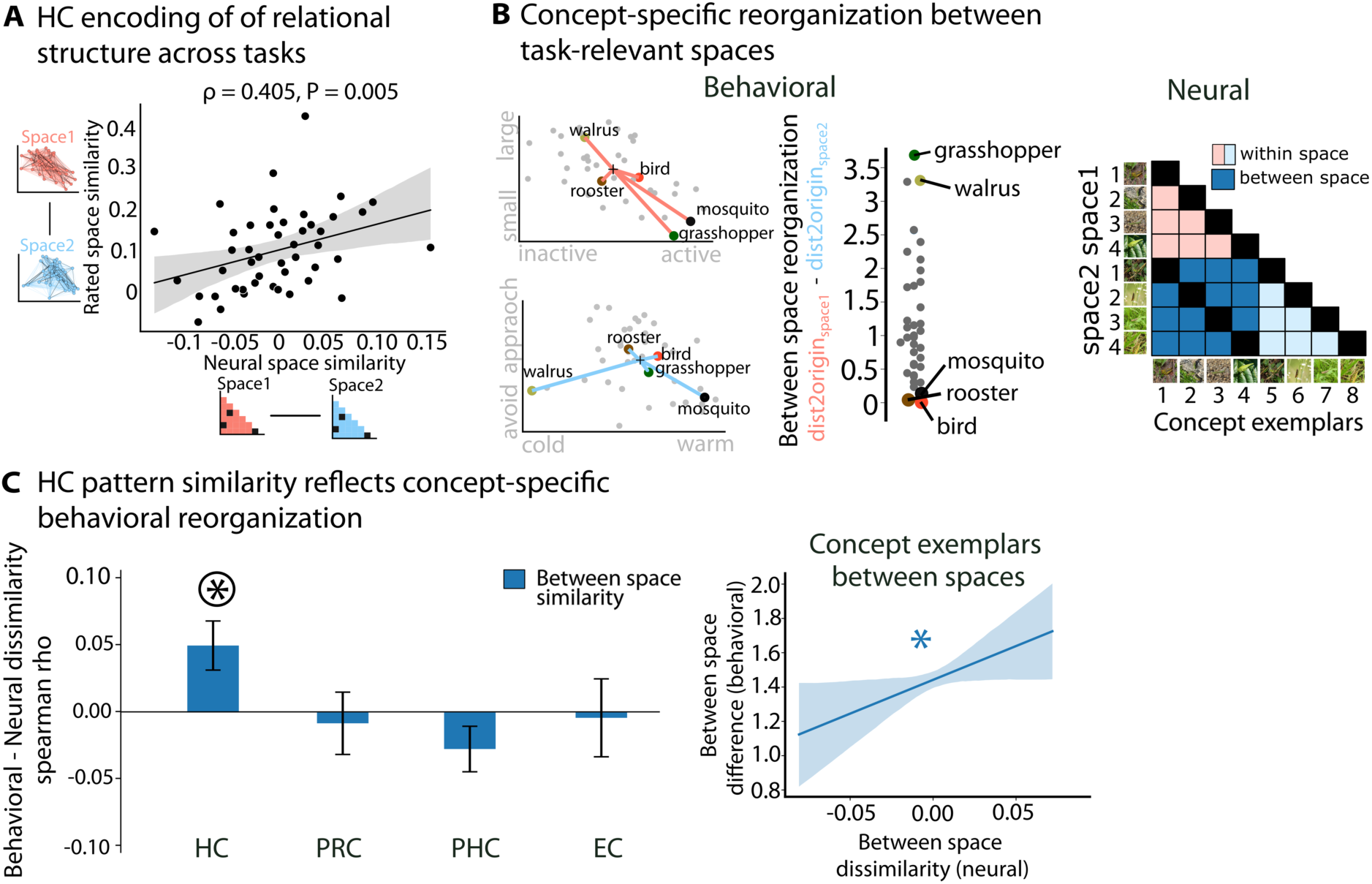
Hippocampal pattern similarity tracks specificity of spaces and context-dependent updating at the level of individual concepts. A) Space-level correlation of behavioral and neural estimates of concept relationships, based only on inferred concept pairs (x-axis: neural space_1_-space_2_ similarity) and behavioral level (y-axis: behavioral space_1_-space_2_ similarity). B) Concept-level reorganization: Schematic depiction of analysis procedure. Participant-specific reconstructed spaces, constituted by the same animal concepts across behavioral contexts (left), and vectors showing the Euclidean distance between exemplary concepts to the origin of the 2D space. Concept-level reorganization across spaces is computed as the difference in the distance of a concept to the origin between the two spaces, i.e. dist2origin_space1_ – dist2origin_space2_ (left). Exemplary distribution of concept reorganization values of one participant (middle). Neural similarity matrix for exemplars of the same animal concept encountered across spaces (right). C) Matching of neural between-space and behavioral concept-level reorganization values, separately for regions of the MTL (left). LMM estimation of neural between-space dissimilarity predicting concept-level remapping. * indicates beta-coefficient with *P* < 0.05 after model comparison. Circled * indicates *P* < 0.05 for one-sample *T* test and line * indicates P < 0.05 for paired *T* test.

Further, we also probed this relationship at the level of individual concepts (i.e., animals), asking whether exemplars of concepts that yield greater reorganization between the two reconstructed representational spaces also show stronger neural differentiation, i.e. lower levels of similarity between exemplars of the same concept that are differentially placed between task-relevant spaces. For example, in one participant, *grasshopper* occupied a relatively peripheral position in the *Size & Activity* space, but not in the *Proximity & Climate* space, indicating behavioral reorganization across spaces (as opposed to concepts that occupy equally peripheral or central positions across spaces; Fig. 4B, left). We quantified this reorganization for each concept by subtracting a concept’s Euclidean distance from the origin ([0,0]) between spaces – a putative concept-specific metric for task-dependent reorganization (Fig. 4B, left) that scales with the differences in concept positions between spaces. Note that the rating-based spaces only contain one representation for each concept, not separate representations for the different exemplars of this concept. On the neural level, we extracted the patterns of each exemplar in each space (four different grasshopper exemplars were presented in each space). We then computed the pairwise neural pattern similarity between all exemplar pairs of the same concept (i.e. all *grasshopper* exemplars) and averaged these values across pairs encountered in the same (within) or different (between) space. This analysis yielded one within-space and one between-space neural similarity measure for each concept (Fig. 4B, right). Finally, we tested whether the behavioral task-dependent reorganization metric was associated with the magnitude of between-space neural pattern dissimilarity across concepts. Positive correlations indicate that concepts whose positions across representational spaces differ more also yield less similar neural patterns between its individual exemplars.

Indeed, we found that correlations were consistently positive across the sample in the HC (*ρ* = 0.05 ±0.12; *T*_46_ = 2.725, *P* = 0.009; Fig. 4C, left). This was not observed in any other MTL subregion (PRC: *T*_46_ = −0.348, *P* = 0.729 | PHC: *T*_46_ = −1.607, *P* = 0.115 | EC: *T*_46_ = −0.138, *P* = 0.891), suggesting a selective role of task-dependent coding in HC. We did not find significant correlations between the reorganization of rating-based distances with neural similarity across exemplars of the same space (i.e. within space similarity; all *P*s > 0.084).

Comparable results were found in an LMM that predicted behavioral concept remapping by fixed effects of within- and between-space HC pattern similarity and participants with random intercepts (Fig. 4C, right): Again, we found that between-space similarity (*Z* = 2.006, *χ*^2^ = 4.02, *P* = 0.045) but not within-space similarity (*Z* = −0.659, *χ*^2^ = 0.19, *P* = 0.659) predicted concept-level reorganization, suggesting that task-specific embeddings in HC are specific to individual concept representations. These findings 1) suggest a selective role for the HC in the embedding of items in task-dependent representational spaces, 2) demonstrate the HCs ability to flexibly reorganize concept relationships to serve behavior in the current task and 3) indicate that tasks requiring concept embeddings with shared features also yield more similar neural representations, suggesting that the HC embeds concepts in abstract spaces that generalize across behavioral contexts.

### Distances in task-relevant representational spaces shape memory formation

Participants provided ratings of the same animal concepts in each block, but each rating was performed on a unique animal exemplar. This allowed us to investigate the formation of episodic memories for these exemplars depending on task-specific relational embeddings in representational spaces. Following fMRI, participants completed a surprise recognition memory task outside of the scanner, where they viewed all previously presented animal exemplars as well as new exemplars of previously presented animal concepts and new animal concepts (Fig. 5A). For each animal image, they indicated on a continuous scale their confidence of having seen it during the rating task; for responses in the “old” range of this scale, they also indicated with a binary response the corresponding representational space (source memory).

**Figure 5:**
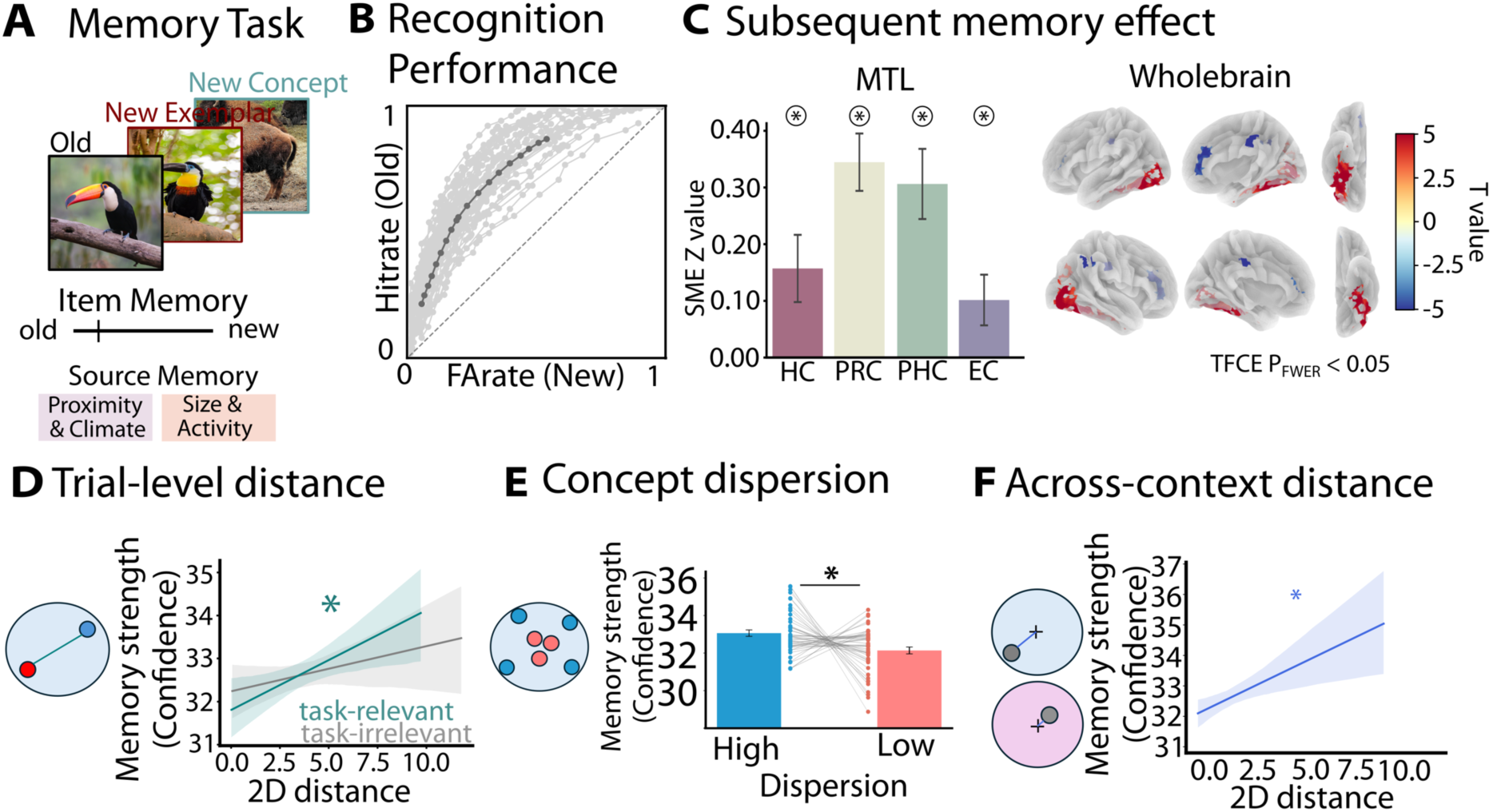
Distance metrics within and across representational spaces are associated with memory strength. A) Recognition memory task. Old: Exemplar images that have been shown during the conceptual navigation task. New exemplar: Novel exemplar of an animal concept that has been shown before. New concept: Animal concept that has not been shown before. B) Receiver-Operator Characteristics (ROC) curve showing the ratio of hits vs. false alarms across the confidence scale (bins of 5 at scale from −50 to +50). Light grey lines represent individual participants; dark grey line indicates sample average. C) Subsequent memory effects in regions of the MTL (left) and across the whole brain (right). D) Trial-level (reference images – blue; comparison images – red), reconstructed (task-relevant – green; task-irrelevant – grey), and concept reorganization (blue) distances from cognitive maps predict memory strength. E) Paired T test of item memory confidence between exemplars of concepts grouped into high and low dispersion, reflecting the relative isolation of a concepts position in task-relevant space. F) Concept-level reorganization indicating the difference in Euclidean distances across spaces as a function of subsequent memory confidence. * indicates significant main effects after model comparison at *P* < 0.05. Circled * indicates *P* < 0.05 for one-sample *T* test. Lined * indicates *P* < 0.05 for paired *T* test.

Participants successfully discriminated between old and new images (*d*-prime: *M* = 1.080, *SD* = 0.256; area under the curve (AUC): *T*_45_ = 29.751, *P* < 0.001; Fig. 5B), and detection of new concepts was better than detection of new exemplars (AUC exemplar vs. concept discrimination: *T*_45_ = −21.414, *P* < 0.001; fig. S6A; for details see Supplement). Source memory accuracy was reliably above chance (i.e., 0.5; *T*_45_ = 4.556, *P* < 0.001), even though it was relatively low overall (0.532±0.047). On the neural level, we tested for subsequent memory effects (SMEs). ROI-based analyses revealed SMEs in all MTL regions (Fig. 5C left; HC: *T*_45_ = 2.639, *P* = 0.012; PRC: *T*_45_ = 6.780, *P* < 0.001; PHC: *T*_45_ = 4.936, *P* < 0.001; EC: *T*_45_ = 2.567, *P* = 0.029). A univariate whole-brain analysis showed positive SME clusters across the VVS (including occipitotemporal, fusiform and parahippocampal cortices; Fig. 3C, right; Table S2), and negative SMEs were found in regions of the default mode network, in middle and anterior cingulate cortex, and in the inferior parietal cortex (Table S2). A parametric modulation analysis confirmed that BOLD activity across MTL and VVS regions increased as a function of memory confidence of subsequently remembered images (fig. S6; Table S2).

We next asked whether distance metrics from individual trials, within task-relevant spaces and across representational spaces were associated with levels of confidence for subsequently remembered images. First, we tested whether task-relevant distances, the major structural property of representational spaces, affected the formation of memories – i.e., whether encoding-related distances associated with each image, relative to the other image presented in the same trial, influenced its memory strength during the subsequent recognition memory test. We set up LMMs predicting memory confidence of subsequently remembered images based on empirically obtained distance ratings during encoding (for details see Methods), and modeled participants with random intercepts. Replicating findings from earlier works (Heinen et al. 2025b), we found a significant main effect of 2D distance (*Z* = 3.552, *χ*^2^ = 12.61, *P* < 0.001), indicating higher memory confidence for images encountered in trials that yielded larger rated Euclidean distances. These effects were observed both for first and second images in each trial (first image: *Z* = 2.342, *χ*^2^ = 5.48, *P* = 0.019; second image: *Z* = 2.765, *χ*^2^ = 7.64, *P* = 0.006). This effect remained when controlling for the cDNN-derived perceptual similarity of both images (for details see Supplement). Further, item memory confidence could not be predicted by the reconstructed task-irrelevant distances (i.e. the distance between both concepts from the subject-specific space that was relevant in other task blocks), as shown in a LMM with reconstructed distances from task-relevant and task-irrelevant distances as predictors of memory confidence (Fig. 5D; task-relevant distances: *Z* = 2.373, *χ*^2^ = 5.63, *P* = 0.0178; task-irrelevant distances: *Z* = 1.094, *χ*^2^ = 1.20, *P* = 0.273). Including the interaction term between task-relevant and task-irrelevant distance did not improve the model (Z = −1.219, *χ*^2^ = 1.48, *P* = 0.223), suggesting that the effect of task-relevant distances on memory confidence is independent of task-irrelevant distance. Finally, the effect of trial-level distance on subsequent item memory confidence remained when including a concept’s distance to the origin in task-relevant space (task-relevant distance: *Z* = 1.963, *χ*^2^ = 3.85, *P* = 0.049; distance to origin: *Z* = 1.033, *χ*^2^ = 1.07, *P* = 0.301). For binary source memory responses, we used a logistic model, which indicated better performance for items encoded with larger task-relevant distances (*Z* = 3.995, *χ*^2^ = 16.01, *P* < 0.001) and even an opposing effect of task-irrelevant distances (*Z* = −2.667, *χ*^2^ = 7.12, *P* = 0.007), suggesting worse source memory if distances in the task-irrelevant spaces were large.

Further, we hypothesized that images would be remembered with higher confidence if their concept occupied a more distinct or isolated position relative to all other concepts in task-relevant space. To quantify this dispersion, we estimated neighborhood density for each concept in task-relevant space and split concepts into groups of high and low dispersion (for details see Methods; fig. S7). Then we tested memory confidence ratings for correctly recognized images across the two dispersion groups. We found that item memory confidence was larger for exemplar images with high (*M*±*SD* = 33.06±1.17) as compared to low (32.14±1.23) dispersion metric (*T*_46_ = 2.365, *P* = 0.011; Fig. 5E), suggesting that a concept’s distinctiveness within task-relevant space is associated with item memory confidence. A LMM confirmed a significant relationship between item memory confidence and rated trial level distance (*Z* = 3.160, *χ*^2^ = 9.98, *P* = 0.001) and a trend for z-scored number of neighbors per concept (i.e. dispersion; *Z* = −1.783, *χ*^2^ = 3.18, *P* = 0.074).

Finally, we tested whether the concept-level reorganization metric, reflecting a concept’s distinctiveness between representational spaces (Fig 4B, left), was associated with item memory confidence. We hypothesized that exemplars of a concept that yield differential distance estimates across spaces would be more confidently remembered as compared to exemplars of a less distinct concept. Indeed, we found that concept-level reorganization predicted higher item memory confidence for Hits (*Z* = 2.758, *χ*^2^ = 7.60, *P* = 0.006). A model with both task-relevant distance and concept remapping showed significant effects of both predictors (task-relevant distance: *Z* = 2.352, *χ*^2^ = 5.53, *P* = 0.019; concept-level reorganization: *Z* = 2.647, *χ*^2^ = 7.01, *P* = 0.008; Fig. 5F).

In sum, while these metrics of concept embeddings within- (trial-level; neighborhood density) and across (distance to origin) task-relevant spaces likely share a common source of variance (for details see Supplement), they support the notion that a concept’s context-derived and behaviorally relevant distinctiveness is associated with successful memory formation – a putative relationship for bridging the gap between relational embeddings in abstract spaces and memory formation associated with the medial temporal lobe.

Contributions of stimulus- and task-dependent representational formats to episodic memories

It is widely assumed that memory traces in the MTL, and in particular the HC, serve as “pointers” or “indices” to neocortical representations (Teyler and DiScenna 1986; Teyler and Rudy 2007). While our findings presented thus far suggest that the accessibility and the representational structure of episodic memory traces are shaped by their embedding in task-relevant representational spaces, the representational formats of neocortical sites in these traces remain unclear.

To address this gap, we identified the neural representations of stimulus-driven perceptual and conceptual formats and compared them to the formats representing task-relevant cognitive maps and to subsequent memory effects. Perceptual formats were quantified via a convolutional DNN model (“AlexNet”) trained to classify images into semantic categories (Krizhevsky et al. 2017). This model hierarchically processes image features of increasing visual complexity. Specifically, we obtained model RSMs for each layer of the DNN (convolutional layers 1-5, fully-connected layers 6-8), representing the similarity structure between all pairs of images, and then used these RSMs to identify brain regions with a similar representational structure (Fig. 6A, left; fig. S8). Replicating findings from earlier work (Cichy et al. 2016; Güclü, and van Gerven 2015; Kuzovkin et al. 2018), we found that regions across occipital cortex and VVS contain representations that correspond to DNN model features, showing matching of basic visual features from convolutional layers (“Conv format”) with representations in occipital cortices, and matching of more complex visual features from fully-connected layers (“Fc format”) with representations in lateral occipital and inferior temporal cortices (Fig. 6A, right; Table S2). Note that for these results, we regressed out correlations with the respective lower-level neural similarity estimates (Fig. 3B-C; Fig. 4C).

**Figure 6:**
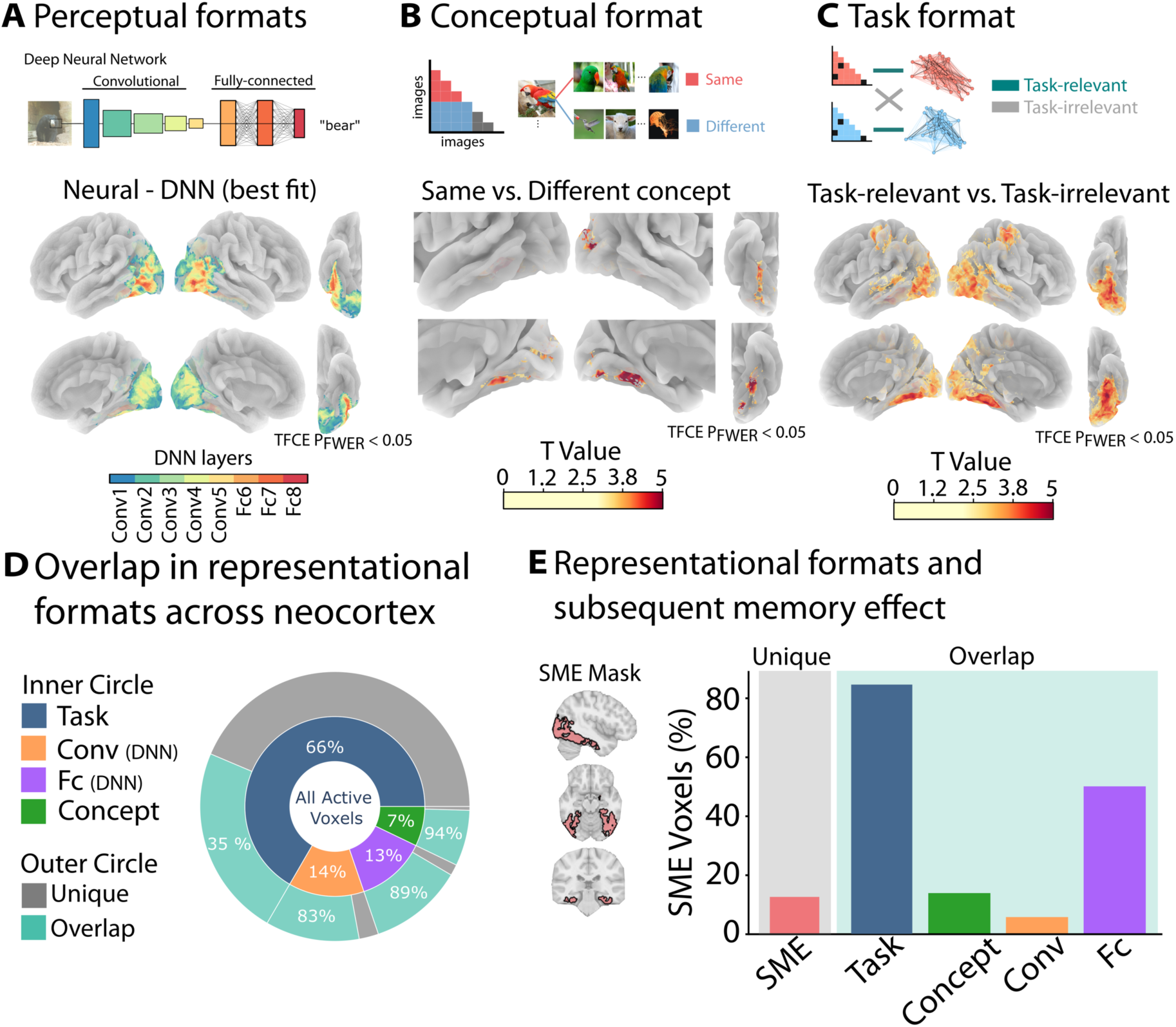
Stimulus-driven and task-dependent representational formats in memory traces. A) Analysis strategy to investigate perceptual formats by comparing pairwise neural similarity matrices with DNN-derived model predictions (top). Visualization of the best layer-model fit plotted onto the surface of a canonical MNI template brain (bottom). All colored voxels contribute to a significant cluster with at least one model layer that survived *P*_FWER_ < 0.05 at the cluster-level after *TFCE*, and the color code depicts the DNN model layer with the best fit for each voxel. B) Analysis strategy to investigate conceptual representational formats across the brain using a searchlight-based RSA approach (top). Concept-specific representations across the VVS (bottom) as indicated by *T* values of significant clusters of the Same vs. Different concept contrast thresholded at the cluster-level with at *P*_FWER_ < 0.05 after *TFCE*. C) Whole brain contrast of task-relevant cognitive map representations (contrast of similarity with task-relevant vs. task-irrelevant maps), *T* values thresholded at the cluster-level with *P*_FWER_ < 0.05 after *TFCE*. D) Distribution of active voxels showing stimulus-driven (perceptual, conceptual) or task-dependent representational formats. Inner circle: Percentage of voxels with distinct representational formats. Outer circle: Percentage of voxels with unique (grey) and overlapping (green) format. E) Distribution of formats across voxels showing positive SMEs as depicted in Fig. 5C.

We next identified stimulus-driven conceptual representations (“Concept format”) by conducting a searchlight-based RSA that compared the neural similarity between exemplars of the same concept vs. different concepts (Fig. 6B, left), while controlling mean BOLD activity and their perceptual similarity (for details see Methods). This analysis yielded medial regions of the VVS extending into bilateral PHC (Fig. 6B, right; Table S3) that are associated with the representation of concepts in a stimulus-driven (i.e., task-independent) format, above and beyond perceptual similarities between concept exemplars. Finally, in comparison to these stimulus-driven (i.e., task-independent) representational formats, we repeated the analysis strategy described in Fig. 3A using a whole-brain searchlight analysis (i.e. “Task-specific format”). We found selective representations of task-relevant vs. task-irrelevant representational spaces across extended areas of ventral and dorsal visual stream (occipitotemporal cortex, fusiform gyrus, middle and superior temporal gyrus, and inferior parietal cortex; Fig. 6C; fig. S9; Table S4), suggesting that task-specific formats extend into occipital and occipitotemporal regions, putatively shaping sensory processing over and above stimulus-driven representational formats, even at early stages of the visual processing hierarchy.

Successful memory may depend on memory traces that comprise multiple representational formats. We therefore sought to determine the relevance of the previously identified formats for the formation of such traces by examining both their spatial overlap with one another and with voxels associated with successful memory encoding (SME). Among all voxels that were involved in the representation of at least one task- or stimulus-dependent format, more voxels reflected task-specific (66%) as compared to stimulus-driven formats (Conv format: 14%; Fc format: 13%; Concept format: 7%; deviation from uniform distribution, binomial test: *P* < 0.001; Fig. 6D, inner circle), suggesting broad spatial tuning towards task-relevant feature representations. This notion is further supported by the finding that most voxels involved in the representation of stimulus-dependent representational formats are non-unique, i.e. are also involved in the representation of other formats (Conv: 83%, Fc: 93%; Concept: 94%). Finally, we quantified the overlap of voxels that were associated with representational formats and successful memory formation as indicated by SME (see Fig. 5C). We observed that 81% of SME voxels overlapped with stimulus-driven and/or task-dependent representational formats, suggesting that memory traces consist of multiple representational formats. Specifically, of these non-unique SME voxels, 84% contributed to a task-specific format (18% of all task-format voxels), 6% to Conv format (6% of all conv-format voxels), 50% to Fc format (58% of all fc-format voxels), and 14% to Concept format (28% of all concept-format voxels; Fig. 6F), suggesting that task-relevant formats may be preferentially but not exclusively encoded into episodic memory.

## Discussion

Representational spaces in the MTL have been demonstrated to support behavior in a variety of tasks that require physical, statistical, social, or abstract reasoning (Constantinescu et al. 2016; Deuker et al. 2016; Garvert et al. 2017; Tavares et al. 2015; Theves et al. 2019). However, most previous studies investigated the emergence of the space itself via repeated exposure or statistical learning, but not on how preexisting conceptual knowledge may be flexibly retrieved and organized in representational spaces in the service of goal-directed behavior. Here, we extend previous findings by demonstrating that similar principles apply to the flexible organization of familiar concepts and arbitrary but task-relevant feature dimensions that can be inferred from prior knowledge. Further, our findings show that this task-specific relational embedding is associated with the formation of novel episodic memories, putatively by reflecting context-dependent embeddings of individual item representations via relational distances within and across representational spaces.

Behaviorally, we found that response times during conceptual comparisons depended on the magnitude of rated distances. Similar effects have been observed in perceptual discrimination tasks and are in line with seminal work suggesting that distances between abstract features may be measured via similarity judgements (Dehaene, Bossini, and Giraux 1993; Hutchinson and Lockhead 1977; Moyer 1973; Moyer and Landauer 1967), putatively reflecting a spatial component in the organization of abstract knowledge (Bottini and Doeller 2020). However, these behavioral observations alone are not sufficient to infer underlying task-specific representational spaces and the pairwise relationships from such spaces may never be actively retrieved during task performance. By only considering distances between untested concept pairs (i.e. inferred pairs) when testing for the neural basis of the underlying representational spaces, we find that concept representations from individual exemplars are indeed embedded in a common space that allows inferences beyond the explicitly probed concepts in each trial.

Our results thus show that MTL organizes concepts in low-dimensional feature spaces, that these maps support task-specific behavior, and that they flexibly update in response to changes in task demands. While we found that neural similarities in MTL reflect task-relevant distances, we also observed negative relationships between neural similarities and task-irrelevant distances in PHC and PRC. Given that natural feature dimensions may not be entirely orthogonal (i.e. larger animals are typically less approachable than smaller animals), negative alignment with task-irrelevant features may reflect a “repulsion” mechanism to enhance the distinctiveness of the representation of task-relevant features (Favila, Chanales, and Kuhl 2016; Chanales et al. 2017).

There has been ongoing debate on whether the MTL is involved in the representation of abstract concepts in a context-dependent or context-independent manner (Quiroga 2020; Suthana et al. 2021). While research on concept cells shows invariant responses to the same concept disregarding contextual aspects (Quiroga 2012; Rey et al. 2015, 2025), other studies show that HC cells rather represent conjunctions of concept- and episodic features (Kolibius et al. 2023) and that they exhibit mixed-selectivity to a variety of features depending on task demands (Donoghue et al. 2023). At the level of macroscopic fMRI responses, we find evidence for context-dependent conceptual representations in HC, MTL and beyond, indicating wide-spread activations in support of task-relevant representational spaces in the brain. Further, context-dependent HC representations of the same animal concept across two behavioral contexts, a putative macroscopic homologue of remapping (Fyhn et al. 2007), argues against context-independent representations in HC and rather suggests concept representations in the MTL that depend on task-demands.

Whole-brain analyses revealed that task-dependent representational formats were also found in ventral temporal and lateral parietal cortices, confirming that representations of task-relevant representational formats are not restricted to the MTL but extend into neocortical regions. Although previous studies identified prefrontal cortex (PFC) in representing spaces of abstract features (Constantinescu et al. 2016) or influencing inference in hippocampal representations (Garvert et al. 2023), we did not find involvement of PFC in representing relational spaces, likely due to the sensory nature of our stimuli and the analysis of task-relevant representational spaces during their viewing. However, while the distinct contributions of MTL and neocortex remain speculative, we found that hippocampal neural pattern similarity was specifically related to both space- and concept-level metrics of concept reorganization, suggesting a proclaimed role of the hippocampal formation in guiding behavioral relevance. Indeed, HC has been described as a hub that integrates perceptual and mnemonic formats (Treder et al. 2021), coordinating representational reinstatement in neocortical areas (Pacheco Estefan et al. 2019). Evidence from rodents further suggests that HC and midbrain regions project context-specific mnemonic information to neocortical layer 1, highlighting task-relevant features, even during early stages of sensory processing (Shin et al. 2021). Our data support this idea by showing that task-relevant information affected neural representations during the viewing of the images, before behavioral ratings were obtained, suggesting that context-specific information interacted with sensory processing of stimuli at multiple stages of the ventral visual stream hierarchy. In the future, studies using invasive electrophysiological recordings in human epilepsy patients, combining high spatial and temporal resolution, could shed further light on the distinct roles of HC and neocortical reinstatement of task-specific formats in humans.

Further, we sought to investigate the functional role of relational item embeddings and the representational formats in the episodic memory traces built from these items. We hypothesized that distances within and across task-relevant representational spaces may index the degree of behavioral and neural discriminability between concepts, providing resistance to interference in neural representations and relating to the strength and accessibility of memories. First, we replicated previous findings by demonstrating that trial-level distances predicted subsequent item memory confidence (Heinen et al. 2025b), suggesting beneficial memory encoding of concept pairs with larger distances in the task-relevant space. This finding is further supported by the notion that exemplar images of concepts with low neighborhood density in task-relevant space (i.e. isolated concepts at the boundaries of concept space distributions) yield higher memory confidence, suggesting that a concept’s relational embedding is associated with subsequent memory. Finally, relational distance effects on memory are supported by our finding that concept-level reorganization was associated with subsequent memory strength as well. While we acknowledge that these distance metrics likely share a common source of variance, combined these findings do suggest a close link between relational distance coding and episodic memory via the embedding of individual items in a spatial coding scheme (Eichenbaum 2017; Whittington et al. 2020), providing evidence for a reconciliation between MTL accounts of spatial navigation and episodic memory via cognitive maps.

We reasoned that memory traces of individual exemplars may consist of both stimulus-driven (i.e. perceptual and/or conceptual) and context-dependent (i.e. task-relevant) representational formats. Replicating previous studies, we observed hierarchically organized perceptual representational formats in occipital cortex and VVS that matched the formats in convolutional and fully-connected DNN layers (Güclü, and van Gerven 2015; Kuzovkin et al. 2018). We (Heinen et al. 2023, 2025a) and others (Davis et al. 2021) have previously shown that either perceptual or conceptual formats can be preferentially encoded and reactivated depending on task demands. Conceptual formats of various object categories have been identified across the neocortex, and animal concepts occurred predominantly in lateral occipital and inferior temporal cortices (Huth et al. 2012). Our results align with these findings as they provide evidence for exemplar-invariant concept representations in VVS. Importantly, compared to these stimulus-driven perceptual and conceptual representations, we observed context-dependent representations in anatomically overlapping brain regions, suggesting multi-layered representational formats and potentially a tuning of stimulus-dependent representations towards task-relevant features in support of behavior. Indeed, our findings suggest that stimulus-dependent conceptual representations as well as task-relevant representations are also found in sensory cortices, suggesting that task-relevant features affect sensory processing even at early stages of the processing hierarchy (Gilbert and Li 2013; Harel, Kravitz, and Baker 2014; Kaiser, Turini, and Cichy 2019). While our findings indicate that representations in feedforward DNNs can account for representations along the visual hierarchy, these models are unable to account for task-dependent representations in sensory cortices. However, recurrent deep neural networks (Kietzmann et al. 2019; Pacheco-Estefan et al. 2024), variational auto encoders (Fayyaz et al. 2022), or deep-Q networks (Walther et al. 2021) could potentially extend the predictive nature of DNN models to account for a broader variety of cognitive functions, behaviorally relevant formats, and their neural representations (Doerig et al. 2023). The relevance of both stimulus- and task-dependent representational formats is further supported by anatomically overlapping effects in brain regions associated with successful memory formation. In these regions, we found that both task-relevant conceptual as well as high-level sensory features were represented, highlighting that both are encoded into memory traces and are relevant to allow for later recognition.

Taken together, our findings demonstrate how natural concepts are embedded in task-relevant cognitive maps to flexibly guide goal-directed decision making. Our results provide insights on how preexisting conceptual knowledge is flexibly accessed and represented and how this relates to the formation of memory traces. We extend previous findings by showing context-dependent hippocampal representations in low-dimensional conceptual spaces of natural stimuli. Further, we show how rated distances from cognitive maps, as well as stimulus- and task-dependent representational formats relate to successful memory formation, suggesting that memory traces comprise multiple distinct representational formats.

## Methods

### Sample

*N* = 53 participants (15 male, 38 female) engaged in the experiment. All participants were right-handed, aged between 18-36 years (24 ± 4 years), without past or current psychiatric disorders and eligible for MRI protocols according to the requirements of Leibniz Institut für Arbeitsforschung (IfADo), Dortmund, Germany. All participants gave written and informed consent and received either monetary compensation (10€ per hour, total 40€) or student credits for their participation. The study was approved by the ethics committee of the Faculty of Psychology at Ruhr University Bochum (proposal #730) in accordance with the declaration of Helsinki.

### Conceptual distance rating task

We designed a conceptual distance estimation task to assess how participants locate and judge animal concepts in low-dimensional but task-relevant representational spaces while undergoing fMRI. On each trial, participants viewed two subsequently presented pictures of animals and were asked to indicate how the second animal related to the first along two predefined feature dimensions. The four conceptual feature dimensions we chose were: *Size* – representing the real-world size of the animals (small - large), *Climate* – indicating the typical climate of the animals’ habitats (cold - warm), *Proximity* – reflecting the distance one would want to have to the animal (avoid - approach), and *Activity* – assessing the perceived activity level of the animal (inactive - active). The pole orientation for each feature dimension (e.g., *Climate*: cold-left, warm-right) was identical for all participants, allowing consistent interpretation of directional ratings.

Across four functional runs, participants completed ratings in two 2D spaces, each defined by a pair of the above feature dimensions (referred to as space1 and space2). Each run consisted of two blocks, one from space1 and one from space2. Across participants, the order of spaces within runs was pseudorandomized (for example, run1: space1 then space2; run2: space2 then space1; run3: space2 then space1; run4: space1 then space2), ensuring that each space occurred equally often as the first and second block across the experiment. The feature dimensions per space did not change for a participant (i.e. for one participant, *Size* would always be paired *Climate* and *Activity* would always be paired with *Proximity*, whereas another participant would e.g. rate *Size* and *Activity*). Across participants eligible for analysis (*N*=46; see exclusion and data quality criteria below), *N*=23 rated *Size* & *Climate* and *Activity* & *Proximity*; *N*=16 rated *Climate* & *Proximity* and *Activity* & *Size*; and *N*=7 rated *Size* & *Proximity* and *Activity* & *Climate*.

At the start of each block, an instruction screen indicated the two relevant feature dimensions for the to be rated space. Each trial consisted of the TR-locked presentation of two animal exemplars (from different concepts) for 1500 ms each, separated by a variable inter-stimulus interval of either two or three TRs (Fig. 1A). Following the images, participants provided two ratings on consecutive screens— one for each dimension relevant to the current block. Within a block, the order of rated dimensions was pseudorandomized at the trial level such that each dimension was presented first in 50% of trials and second in the other 50%. Participants responded using two 5-button boxes, one held in each hand, which mapped onto a 10-point Likert scale ranging from −5 (left little finger) to +5 (right little finger), allowing participants to indicate the direction and magnitude of the relationship between the second and the first animal along each dimension with respect to the fixed left–right poles. Each rating screen remained visible until a response was provided or for up to a maximum of 8000 ms. Missed responses were followed by an instruction screen to please respond faster in the next trial. On average, participants missed 0.5 ± 0.65 ratings. These trials (image and rating events) were excluded from all analyses.

For each participant, the stimulus set comprised 40 animal concepts randomly drawn from a larger pool of 93 concepts (see below). The same 40 concepts were used in both spaces throughout the experiment. Participants completed eight task blocks in total (4 runs à 2 spaces), and each concept appeared exactly once in every space. Thus, each concept was shown eight times across the experiment—four times in space1 and four times in space2 (once per space)—ensuring equal numbers of appearances in both spaces. For each appearance of an animal concept, a unique exemplar image was used, such that no image was ever repeated; only the concept recurred across spaces. In total, this yielded 320 unique images (8 exemplars × 40 concepts) and enabled subsequent tests of episodic, image-specific memory in a follow-up recognition task.

Each block consisted of 20 trials. Each trial presented two different concepts (one image per concept), so that the 40 concepts were covered exactly once per block. Across the eight blocks, participants therefore completed 160 trials in total. Within blocks each concept appeared once and was never paired with itself; pairings were re-randomized across blocks and across spaces to produce largely non-overlapping sets of pairs.

### Recognition memory task

The recognition memory task was performed on a laptop outside of the MRI scanner. In total, participants saw 480 unique animal exemplar images split into 6 blocks of 80 trials. Of those 480 images, 320 were exemplar images from the first task (i.e. old); 80 images were new exemplars (i.e. there were 2 new exemplars of each of the 40 animal concepts encountered in the first task); and 80 images were new concepts (2 exemplars of 40 novel animal concepts; Fig. 1D).

Each trial began with the presentation of a fixation cross for 500ms. This was followed by the presentation of the animal image which was shown until a response was given. Below the image, participants saw a continuous rating slider that ranged from “old” (−50) to “new” (+50) in steps of 1. Participants were asked to indicate their confidence of having seen the presented image (the exact exemplar, not the animal concept) in the conceptual navigation task using the continuous slider scale. They were explicitly instructed to not only give binary judgements (old, new) but to indicate their level of confidence using the entire rating scale. In case participants indicated having seen the presented image (i.e. responded “old” by selecting a point on the left side of the scale), they were probed on source memory. In a forced choice task, they indicated the space in which they had encountered the presented image (e.g. *Proximity* & *Climate* or *Size* & *Activity*). In case participants indicated that the presented image was new, no source memory test was performed and the experiment continued with the next trial. As a measure of recognition performance, we computed *d*-prime (Yonelinas 2002; Yonelinas and Parks 2007) defined as the standardized difference of hits vs. false alarms to control for a participant’s response bias. Further, we tested whether the area under the curve (AUC) across confidence bins (bin size = 5) was reliably above change (0.5) using one-sample *T* tests.

### Stimuli & Software

All animal images were taken from the *THINGS* database (Hebart et al. 2019) which provides multiple high-resolution natural images of various different animal concepts. We excluded concepts if they were highly-arousing (e.g. spiders, snakes, ticks) or if not enough images with single animal exemplars were available. We included 93 animal concepts, with ten suitable exemplar images per concept (eight for the conceptual navigation task and two novel exemplars for the recognition memory test). For each participant, we randomly selected 40 animal concepts for the conceptual navigation task, and another 40 animal concepts during the recognition memory task. The experimental tasks were programmed in jspsych (Leeuw, Gilbert, and Luchterhandt 2023) and run in a browser on a Windows 10 Desktop Computer (conceptual navigation task) and a laptop (recognition memory task), respectively. Image presentations were locked to the onset of a TR, realized via simulated button presses emulated by the MR scanner and recognized by the experiment. Exemplary animal images in the figures of the manuscript are either license-free versions (Creative Commons Zero; CC0) provided by the THINGS+ database (Stoinski, Perkuhn, and Hebart 2022) or custom photographs.

### Rated and reconstructed Euclidean distances in conceptual spaces

During the conceptual distance rating task, we obtained 80 pairwise similarity ratings for each conceptual feature dimension. We used these ratings to compute a trial-level Euclidean distance metric indicating the relationship of each pair of animals given the task-relevant features. To do so, we conceptualized that the relative position of the animals in two-dimensional feature space where the x- and y-coordinates correspond to the first and second rating response, respectively (Fig. 2A). As participants rated the second animal relative to the first, we defined the position of the reference image (first image) as the origin at coordinates (0,0) and the position of the comparison image (second image) at coordinates (rating1, rating2) to compute their Euclidean distance. This yielded a trial-level Euclidean distance metric derived from behavioral ratings given two conceptual features.

However, this trial-level Euclidean distance metric is restricted to pairs of animal concepts that were explicitly rated during the experiment, comprising only a fraction (∼10 %) of all possible pairwise combinations of the 40 animal concepts (40×39/2 = 780 unique concept pairs for 40 animal concepts). To estimate conceptual feature distances of all concept pairs, we approximated the optimal position for each concept and feature dimension to reconstruct Euclidean distance metrics between all pairs of animal concepts. To do so, we set up a compressed sparse column matrix (design matrix M) of shape (#ratings x #animals; 80 x 40) for each dimension, where each row in M is composed of {-1,0,1} at cells indexing those animal concepts that were explicitly rated in a given trial, with −1 indicating the cell of the reference image and +1 indicating the cell of comparison image (Fig. 2B). Then, separately for each feature dimension (i.e. *Size*, *Climate*, *Proximity*, *Activity*), we use a least-squares approach to solve M to best approximate the behavioral ratings obtained on each dimension, yielding a solution for each animal reflecting its estimated position on the feature dimension given the entirety of ratings obtained by a participant. The estimated positions for each animal concept were then used as x- and y-coordinates in the two participant-specific spaces (Fig. 2C). Importantly, we estimated both participant-specific reconstructions of spaces (i.e. based on ratings obtained by a participant) as well as global reconstructions of spaces (i.e. based on all ratings obtained across all participants). For global space reconstructions, we combined individual dimensions into all possible 2D spaces (i.e. *Activity* & *Size*, *Activity* & *Climate*, *Activity* & *Proximity*, *Size* & *Climate*, *Size* & *Proximity* and *Activity* & *Proximity*). Further, we quantified whether reconstructed spaces indeed differed between feature dimensions, i.e. whether space reconstructions were specific to their conceptual feature dimensions. Therefore, we tested across participants, whether reconstructed spaces of the same dimensions were more similar as compared to reconstructions of spaces with different dimensions. Specifically, using Spearman’s rank correlation, we computed the similarity between participant-specific and group-level space reconstructions, where for example, a participant-specific *Activity & Climate* space was correlated with the groups *Activity & Climate* space (same space) as well as with the groups *Size & Proximity* space (different space). Note that the group space reconstructions were created for each participant individually, based on all other participants to avoid circular comparisons to a group that contains the same participant.

### Validation of space reconstructions

To ensure that the obtained solutions for each dimension fit the behavioral responses better than chance, we compared the error in the empirical least-squares solution to a null distribution of random data. Specifically, we estimated concept positions in each participant, extracted the least squares error from the empirical estimation and repeated the analysis 5,000 times, each time shuffling the behavioral responses within participant while keeping the design matrix M constant. Then, we averaged the empirical error across participants and ranked the group-mean in the averaged permutation distribution to obtain a *P*-value of the empirical value. To ensure that reconstructed spaces meaningfully generalize to untested concept pairs, we additionally performed a leave-one-pair-out cross-validation for each participant and rated feature dimension. Specifically, we left out each explicitly rated concept pair, estimated the participant-specific model based on the remaining responses, correlated between empirically obtained and model predicted ratings using Spearman’s rank correlation, and tested whether obtained correlations were substantially different from zero across the sample.

### Linear Mixed Models

We performed linear mixed models of the statsmodels package (Perktold, J. et al. 2023) in Python 3.8. To determine the influence of different fixed effects on behavioral metrics, we set up a full model containing all fixed effects and participants as a random effect. We then use a likelihood test to determine whether the exclusion of the factor of interest significantly reduces the explained variance compared with the full model. For main effects with one predictor the reduced model included a constant as fixed effect (1). For main effects with two or more predictors the reduced model contained all other main effects except the effect of interest. For interaction effects, the reduced model included both main effects without the interaction term. For each fixed effect, we report *Z*-statistics from the initial estimation of the model and the respective likelihood derived from model comparison (χ2) and its significance level (*P*).

### fMRI acquisition

Imaging data were collected on a 3T Siemens Prisma scanner (Magnetom) at the IfADo, Dortmund, Germany with a 64-channel head coil. We acquired four task-based functional T2* weighted runs using echo-planar imaging (EPI) sequence at multiband acceleration factor 6 (isotropic voxel resolution = 1.6 mm, FOV = 132×132×78, FA = 52°, TR = 1200ms, TE = 31.4ms, phase-encoding direction PE = A-P). The average number of volumes per functional run was 514 ± 47 volumes, with a total average of 2,059 ± 170 volumes per participant. Before the first and after the last functional runs, we acquired 5 volumes with a reversed PE EPI fieldmap sequence (PE = P-A) to correct for susceptibility-induced distortions in the functional images. The acquisition of each EPI scan was preceded by the acquisition of a single-band reference volume. Next, we acquired two T1-weighted anatomical scans, using MPRAGE (isotropic voxel resolution = 1mm, FOV = 256×256×176, FA= 7°, TR = 2530ms, TE = 2.36ms) and MP2RAGE sequences (isotropic voxel resolution = 1mm^3^, FOV = 256×240×176, FA_1/2_ = 4°/5°, TI_1/2_ = 700ms/2500ms, TR = 5000ms, TE = 2.03ms). Finally, we acquired a T2-weighted structural scan using turbo spin-echo sequence (isotropic voxel resolution = 1 mm^3^, FOV = 256×256×224, FA = 120°, TR = 2800ms, TE = 405ms). As T1-weighted reference image we used the MP2RAGE scan, that was preprocessed using *presurfer (*freely available at https://github.com/srikash/presurfer)(Kashyap 2021) for compatibility with freesurfer-based reconstruction. For one subject, we used the MPRAGE anatomical image due to movement distortions during the acquisition of the MP2RAGE sequence. Raw MRI data were organized according to “Brain Imaging Data Structure” (BIDS) (Gorgolewski et al. 2016) using *heudiconv* (v. 0.13.1 freely available at: https://github.com/ReproNim/reproin) and preprocessed using the *fMRIPrep* (Esteban et al. 2019) toolbox (v. 21.0.2) run inside a Docker (Merkel 2014) container. The following sections (Preprocessing of B0 inhomogeneity mappings, Anatomical data preprocessing, Functional data preprocessing) are boilerplate text generated by *fMRIPrep*.

#### Preprocessing of B0 inhomogeneity mappings

A total of 4 fieldmaps were found available within the input BIDS structure. A B0-nonuniformity map (or fieldmap) was estimated based on two (or more) echo-planar imaging (EPI) references with topup (Andersson, Skare, and Ashburner 2003); FSL 6.0.5.1:57b01774).

#### Anatomical data preprocessing

A total of 1 T1-weighted (T1w) images were found within the input BIDS dataset. The T1-weighted (T1w) image was corrected for intensity non-uniformity (INU) with N4BiasFieldCorrection (Tustison et al. 2010), distributed with ANTs 2.3.3 (Avants et al. 2008) (RRID:SCR_004757), and used as T1w-reference throughout the workflow. The T1w-reference was then skull-stripped with a Nipype implementation of the antsBrainExtraction.sh workflow (from ANTs), using OASIS30ANTs as target template. Brain tissue segmentation of cerebrospinal fluid (CSF), white-matter (WM) and gray-matter (GM) was performed on the brain-extracted T1w using fast (Zhang, Brady, and Smith 2001) (FSL 6.0.5.1:57b01774, RRID:SCR_002823). Brain surfaces were reconstructed using recon-all (Dale, Fischl, and Sereno 1999) (FreeSurfer 6.0.1, RRID:SCR_001847), and the brain mask estimated previously was refined with a custom variation of the method to reconcile ANTs-derived and FreeSurfer-derived segmentations of the cortical gray-matter of Mindboggle (Klein et al. 2017) (RRID:SCR_002438). Volume-based spatial normalization to one standard space (MNI152NLin2009cAsym) was performed through nonlinear registration with *antsRegistration* (ANTs 2.3.3), using brain-extracted versions of both T1w reference and the T1w template. The following template was selected for spatial normalization: ICBM 152 Nonlinear Asymmetrical template version 2009c (Fonov et al. 2009) (RRID:SCR_008796; TemplateFlow ID: MNI152NLin2009cAsym).

#### Functional data preprocessing

For each of the 4 BOLD runs found per subject (across all tasks and sessions), the following preprocessing was performed. First, a reference volume and its skull-stripped version were generated by aligning and averaging 1 single-band reference (SBRef). Head-motion parameters with respect to the BOLD reference (transformation matrices, and six corresponding rotation and translation parameters) are were estimated before any spatiotemporal filtering using mcflirt (Jenkinson et al. 2002) (FSL 6.0.5.1:57b01774). The estimated fieldmap was then aligned with rigid-registration to the target EPI (echo-planar imaging) reference run. The field coefficients were mapped on to the reference EPI using the transform. BOLD runs were slice-time corrected to 0.545s (0.5 of slice acquisition range 0s-1.09s) using 3dTshift from AFNI (Cox and Hyde 1997) (RRID:SCR_005927). The BOLD reference was then co-registered to the T1w reference using bbregister (FreeSurfer) which implements boundary-based registration (Greve and Fischl 2009). Co-registration was configured with six degrees of freedom. First, a reference volume and its skull-stripped version were generated using a custom methodology of fMRIPrep. Several confounding time-series were calculated based on the preprocessed BOLD: framewise displacement (FD), DVARS and three region-wise global signals. FD was computed using two formulations following (Power et al. 2014) (absolute sum of relative motions) and (Jenkinson et al. 2002) (relative root mean square displacement between affines). FD and DVARS were calculated for each functional run, both using their implementations in Nipype (following the definitions by (Power et al. 2014)). The three global signals were extracted within the CSF, the WM, and the whole-brain masks. Additionally, a set of physiological regressors were extracted to allow for component-based noise correction (Behzadi et al. 2007) (CompCor). Principal components were estimated after high-pass filtering the preprocessed BOLD time-series (using a discrete cosine filter with 128s cut-off) for the two CompCor variants: temporal (tCompCor) and anatomical (aCompCor). tCompCor components were then calculated from the top 2% variable voxels within the brain mask. For aCompCor, three probabilistic masks (CSF, WM and combined CSF+WM) were generated in anatomical space. The implementation differs from that of Behzadi et al. in that instead of eroding the masks by 2 pixels on BOLD space, the aCompCor masks were subtracted a mask of pixels that likely contain a volume fraction of GM. This mask was obtained by dilating a GM mask extracted from the FreeSurfer’s aseg segmentation, and it ensures components are not extracted from voxels containing a minimal fraction of GM. Finally, these masks were resampled into BOLD space and binarized by thresholding at 0.99 (as in the original implementation). Components were also calculated separately within the WM and CSF masks. For each CompCor decomposition, the k components with the largest singular values were retained, such that the retained components’ time series are sufficient to explain 50 percent of variance across the nuisance mask (CSF, WM, combined, or temporal). The remaining components were dropped from consideration. The head-motion estimates calculated in the correction step were also placed within the corresponding confounds file. The confound time series derived from head motion estimates and global signals were expanded with the inclusion of temporal derivatives and quadratic terms for each (Satterthwaite et al. 2013). Frames that exceeded a threshold of 0.5 mm FD or 1.5 standardised DVARS were annotated as motion outliers. The BOLD time-series were resampled into standard space, generating a preprocessed BOLD run in MNI152NLin2009cAsym space. First, a reference volume and its skull-stripped version were generated using a custom methodology of *fMRIPrep*. All resamplings can be performed with a single interpolation step by composing all the pertinent transformations (i.e. head-motion transform matrices, susceptibility distortion correction when available, and co-registrations to anatomical and output spaces). Gridded (volumetric) resamplings were performed using antsApplyTransforms (ANTs), configured with Lanczos interpolation to minimize the smoothing effects of other kernels (Lanczos 1964). Non-gridded (surface) resamplings were performed using mri_vol2surf (FreeSurfer).

### fMRI data quality control and exclusion criteria

As a decision criterion we required average framewise displacement (Power et al. 2012) (FD), quantifying estimated bulk-head motion, to be lower than 2mm across runs and no single displacement to be larger than 3.9mm. *N*=3 participants did not meet these criteria and were removed from all functional and behavioral analyses. The remaining participants had an average FD of 0.156 ± 0.015 mm across runs. We excluded another *n* = 3 participants because of a zipper artifact contaminating more than two functional runs of functional T2* weighted data runs, likely caused due to a malfunctioning head-coil. The zipper artefact was also present in *n*=8 participants in one half of one functional run. We did not exclude these participants entirely but merely excluded the data from the contaminated runs from functional analyses and confirmed that the remaining data was non-contaminated and of sufficient quality. Importantly, as described in earlier sections, all experimental conditions, the number of concept presentations and ratings per dimension were equally distributed across functional runs. Therefore, for affected participants, the number of trials per condition were still constant, merely reducing to 60 instead of 80 trials per condition, ensuring balanced contrasts estimated at the subject-level. Finally, one participant aborted data acquisition and had to be excluded from further analyses, resulting in a final sample of *N*=46 participants.

To ensure high data quality, raw and preprocessed anatomical and functional scans were visually inspected by trained experts and evaluated by computing metrics for quality control using the LN_SKEW function (v. 2.5.2; https://github.com/layerfMRI/LAYNII) provided by LayNii (Huber et al. 2021). Finally, personal facial features were removed from anatomical T1w scans using pydeface (v. 2.0.0; https://github.com/poldracklab/pydeface/blob/master/Dockerfile) (Gulban et al. 2022).

### Regions of interest

For wholebrain searchlight analyses, we restricted the search space to subject-specific masks of gray matter obtained from *freesurfer* segmentation. Since we had strong hypotheses for effects in the MTL, and more specifically HC, we additionally conducted analyses in predefined regions of interest (ROI) in PHC, PRC, EC which were obtained from detailed participant-specific segmentations in native space using the cloud-based Automated Segmentation of Hippocampal Subfields (ASHS) ASHS-PMC-T1 atlas (Yushkevich et al. 2015; Xie et al. 2016). For descriptive statistics on average number of voxels and temporal signal-to-noise ratio please see Supplement (fig. S10). For second level analyses, all participant-specific masks were transformed to MNI space (template MNI152NLin2009cAsym) using functionality provided by ANTs and merged into one group-level mask.

### Univariate fMRI analyses

We identified regions across the brain where BOLD activity during viewing of images in the conceptual distance rating task was associated with later memory. The MNI-transformed data for each functional run was modeled using a general linear model (GLM) composed of stimulus onset regressors for 1) subsequently remembered images, 2) subsequently forgotten images, 3) rating events, 4) button presses, and 5) a parametric modulator weighting the magnitude of subsequently remembered images by the mean-centered memory confidence ratings obtained during the recognition task. The modulated regressor followed the unmodulated image onset regressor for remembered scenes in the model (Mumford, Poline, and Poldrack 2015). The duration of image events as well as their parametric modulation was equal to the presentation duration of images, i.e. 1500ms. In the GLM, we included six motion regressors (three rotational, three translational), six drift model regressors, and an intercept as regressors of no interest. In case *fMRIPrep* confounds estimation indicated the presence of non-steady-state volumes in the beginning of a functional run, we included those in the model, with separate onset regressors for each marked non-steady state volume. Each regressor of interest was convolved with a double-gamma hemodynamic response function (HRF). On the subject-level, we applied 4.8mm smoothing and used *Z*-scored beta maps for the individual regressors of interest for subsequent second-level analysis.

### Extraction of beta-series

To estimate event-specific BOLD activation maps, we followed the least-squares separate (LS-S) approach for subsequent multivariate pattern analyses, which has been described to be well suited for fast event-related designs (Abdulrahman and Henson 2016; Mumford et al. 2012). Separately for each functional run and event (reference image, comparison image, rating1, rating2) we set up a GLM with an event onset regressor containing only the onset of the to-be-estimated event of interest and two other regressors containing all event onsets of the remaining image and rating events. Durations for image events were equal to the stimulus presentation (1.5s) while the duration of rating events corresponded to the response times associated with each rating event. Additional confound regressors were six regressors modelling motion (three translational, three rotational), six drift-model regressors, and one constant intercept regressor. In case *fMRIPrep* confounds estimation indicated the presence of non-steady-state volumes in the beginning of a functional run, we included those in the model, with separate onset regressors for each marked non-steady state volume. After model estimation, we extracted the beta-values for the event of interest and used the resulting brain activation map for subsequent multivariate searchlight representational similarity analysis.

### Representational similarity analysis

We conducted representational similarity analysis based on the event-specific activity maps obtained from LS-S trial-level beta estimation (Mumford et al. 2012). The following procedure was equally applied to predefined regions of interest in MTL or individual searchlight spheres in native space. For all individual presentations of an animal image (only image events), we computed pairwise correlations across the multi-voxel activity patterns using Spearman’s rank correlation, resulting in a symmetric image x image similarity matrix (size = 320×320) for each region of interest or searchlight that was Fisher-*Z*-transformed before further computations. Importantly, to minimize the influence of false positives in pattern similarity estimates, we restricted all analyses to between-run similarities only (Mumford, Davis, and Poldrack 2014), thereby avoiding confounds due to temporal autocorrelations among volumes acquired in the same functional run.

### Searchlight

We conducted wholebrain RSA (Kriegeskorte, Goebel, and Bandettini 2006) with the searchlight approach, building on python-based functionality provided by the *rsatoolbox* (Nili et al. 2014) (v 0.1.5). We scanned the gray-matter masks obtained from freesurfer-based registration in native space of each participant with spherical searchlights of radius n = 5 voxels and required a minimum of 50 voxels per searchlight to be considered for further analyses. For each individual searchlight sphere, we controlled for mean BOLD activity and visual similarity (for details see below). The resulting first-level maps after RSA contrasts were then transformed into MNI space using ANTs (Avants et al. 2014) *applyTransforms* (v 2.5.0) for second-level analysis.

### Second-level analysis

First-level statistical maps from univariate and searchlight RSA analyses were subjected to non-parametric second-level analysis with 5,000 permutations and threshold-free cluster enhancement (Smith and Nichols 2009) (*TFCE*) as implemented in *nilearn*. We report *T* and *TFCE* statistics for clusters surviving *TFCE*-correction for multiple comparisons at a family-wise error rate (*FWER*) of *P*_FWER_<0.05. MNI coordinates, and peak statistical values associated with each cluster as well as associated anatomical labels for peak voxels according to the AAL2 (Rolls, Joliot, and Tzourio-Mazoyer 2015) or ASHS atlas were extracted using AtlasReader (Notter et al. 2019) and are reported in the Supplement.

### Deep neural network model

We used a pre-trained convolutional deep neural network (cDNN), “AlexNet”(Krizhevsky et al. 2017), as implemented in the Caffee framework (Jia et al. 2014) in python 2.7. To analyze cDNN similarities, we first generated features for each layer of the AlexNet for all images. In each layer, we averaged across the spatial dimension keeping only one value per feature. We then correlated the features of each image with all other images using Spearman’s rank correlation, resulting in 8 image x image correlation matrices, one for each layer. These correlation maps represent the similarity of each image to all other images across the layers of the cDNN based on perceptual properties of increasing complexity, starting with similarities based on early visual information in convolutional layers and concluding with higher-order visual information in fully-connected layers.

To relate the representational formats of the cDNN layers to brain representations, we conducted searchlight-based RSA separately for each layer of the network. For each searchlight, we took the fisher-*Z*-transformed similarities between all images (image x image matrix) and used Spearman’s rank correlation to compare matrices between brain and model layers, yielding a correlation map across the brain for each participant. To determine the best layer fit for each voxel in the brain (Fig. 6B) we first identified all voxels that were part of at least one layer-specific cluster after *TFCE* correction at *P*_FWER_ < 0.05. If a voxel showed significant alignment with multiple layers of the cDNN, we compared the voxel-level *T*-value from second-level comparisons and color coded the voxel by the layer with the highest *T*-value contrast.

### Controlling neural similarity for mean BOLD activity and perceptual formats

We regressed variance from neural similarity estimates attributable to 1) the mean levels of BOLD activation during image presentation (Davis et al. 2014; Mumford et al. 2012) and 2) the similarity attributable to visual features as predicted by the cDNN. For each ROI or searchlight sphere, we set up a GLM predicting the neural pattern similarity for all between-run similarities by two regressors corresponding to the mean BOLD activation across voxels for each image that contribute to the pairwise similarity estimate and 2) eight regressors corresponding to the perceptual similarity of individual cDNN layers (conv1-5, fc6-8) reflecting the magnitude of perceptual similarity of increasingly complex visual formats (fig. S3; for similar procedure see (Kaiser et al. 2019)). The obtained residuals from this GLM were used for further analyses. Note that to identify cDNN-predicted perceptual formats of the brain, we only regressed mean BOLD activity from neural similarity estimates. To correct for multiple comparisons across layers, we ensured that all *P-*values of detected clusters survive Bonferroni-correction

### Stimulus-dependent conceptual representations

We conducted a wholebrain searchlight analysis testing for stimulus-driven conceptual representations in the brain. To do so, we used neural similarity estimates adjusted for mean BOLD activity and visual formats (see above) for all images. Then, we average across all pairwise similarities of animal exemplars belonging to the same animal concept (e.g. bird1 – bird2, bird2 - bird3) and contrasted it with the average pairwise similarity between animal exemplars belonging to a different animal concept (e.g. bird 1 – monkey 1, bird 1 – sheep 2).

### Testing task-specific representational spaces via alignment of neural and reconstructed concept relationships

To compare neural representational similarities with reconstructed distances from behavioral cognitive maps we used the residuals after regression of BOLD magnitude and DNN-predicted perceptual similarities of all between-run similarities of the image x image representational similarity matrix (RSM), separately for space1 and space2. Next, for each space, we averaged similarities across exemplars of the same animal, yielding a space-specific RSM of shape concept x concept. Similarly, we obtained a dissimilarity matrix (RDM) of behavioral distance for pairs of concepts by extracting the task-relevant (same space) and task-irrelevant (difference space) distances from reconstructed cognitive maps. Importantly, to allow direct comparisons between inferred concept relationships, we used the subset of concept pairs that were inferred in both spaces. Therefore, in each subject, we determined pairs of concepts that were explicitly probed during distance ratings and excluded these from both neuronal and rated matrices. Next, we correlated vectors of neural and behaviorally reconstructed concept pairs, separately for space1 and space2, with matching order (i.e. task-relevant space; neural_space1_ – reconstructed_space1_; neural_space2_ – reconstructed_space2_) and with non-matching order (i.e. task-irrelevant space: neural_space1_ – reconstructed_space2_; neural_space2_ – reconstructed_space1_) and averaged task-relevant and task-irrelevant maps across spaces. Finally, we subtracted task-irrelevant from task-relevant correlations to obtain a map of task-specific correlations. Since we correlated neural similarities with rated distances, we sign-flipped the resulting correlation values for further comparisons. This analysis yielded three statistical maps (task-relevant; task-irrelevant; task-specific) that were used for second level analysis.

### Space- and concept-level reorganization

To test the relatedness of reconstructed and neural space representations, we conducted a correlation analysis using Spearman’s rank correlation between rated and neural RSMs (i.e. neural space_1_ – neural space_2_; rated space_1_ – rated space_2_). For neural RSMs, we used the concept x concept RSMs computed across voxels in predefined ROIs of the MTL. This yielded two correlation values per participant, i.e. space_1_-space_2_ similarity based on reconstructed and neural representational spaces. In a second step, we correlated between neural and rated space_1_-space_2_ similarity values across participants.

Further, we quantified behavioral between-space reorganization of individual concepts between both representational spaces rated by a participant. Specifically, we capitalized on participant-specific reconstructed representational spaces and for each concept extracted the Euclidean distance to the origin within each space (origin [0,0]), yielding two distance metrics per concept. Note that we used the distance to the origin since this metric is agnostic to the direction of the vector to the origin which is crucially different between both spaces that are constituted of different conceptual features that are not directly comparable. Finally, for each concept, we computed the difference in the distance to the origin between spaces (i.e. concept remapping = dist2origin_space1_ – dist2origin_space2_) yielding one across-space reorganization value per animal concept. Note that this metric quantified the degree to which a concept is differently positioned across representational spaces. Specifically, this metric would be low for animals equally exposed position in both maps (i.e. a lion may be considered both big, avoidable, active and living in a warm climate), but high for animals that are attributed special characteristics in one but not the other space (i.e. a grasshopper may be considered relatively small and active but indifferently rated regarding *Proximity* or *Climate*). Note that we chose this metric, as it is agnostic to the directionality and pole orientations of feature dimension axes.

Separately for each MTL region, we again extracted pairwise similarities across all images and averaged across all exemplars of the same concept that were encountered in the same space or in a different space, yielding two similarity estimates per concept. Next, within participants, we correlated across concept-level reorganization values and concept averaged within and between space neural similarity using Spearman’s rank correlation, flipped the sign such that behavioral across-space distance and neural similarity share directionality (i.e. distance – dissimilarity) and performed a one-sample *T* test across subjects. Accordingly, we conducted an LMM, predicting behavioral concept reorganization with same and different neural similarity as fixed effects and participants as random effects.

## Author Contributions

Conceptualization: E.M.B.R, R.H., N.A.H., E.Z.P., M.K., N.A.

Methodology: E.M.B.R., R.H., N.A.H., E.Z.P., M.K., K.A., N.A.

Software: E.M.B.R, R.H., N.A.H., M.K., K.A.

Validation: E.M.B.R, R.H., M.K., K.A.

Formal Analysis: E.M.B.R., R.H., M.K.

Investigation: E.M.B.R., R.H., N.A.H.

Resources: E.M.B.R., R.H., N.A.H., K.A.

Data curation: E.M.B.R., R.H., N.A.H.

Writing – original draft: E.M.B.R.

Writing – review & editing: E.M.B.R., R.H., E.Z.P., M.K., K.A., N.A.

Visualization: E.M.B.R., R.H., N.A.H.

Supervision: N.A.

Project administration: E.M.B.R., N.A.

Funding acquisition: N.A.

We thank E. Genç and M. Burke for their contributions to MRI sequence development and technical support during data acquisition. Further, we thank L. Ehrlich, A. Venjakob and R. Merz for assistance during data collection and preparation.

Calculations (or parts of them) for this publication were performed on the HPC cluster Elysium of the Ruhr University Bochum, subsidised by the DFG (INST 213/1055-1).

## Data and materials availability statement

All original code, behavioral data, de-identified anatomical and processed functional MRI data have been deposited at Zenodo and will be publicly available as of the date of publication in a peer-reviewed journal.

## Supplemental information

Supplementary Text

Supplementary Figures (S1–S10)

Supplementary Tables (T1-T5)

## Supplementary Materials

### Supplementary Text

#### Stimulus-dependent perceptual feature distance does not predict rating RTs and item-memory confidence

Besides task-relevant feature distance ratings, we tested whether response times during ratings in individual trials were influenced by perceptual similarities between images of a trial. We used a feedforward convolutional neural network with five convolutional and three fully-connected layers (“AlexNet”; Krizhevsky et al. 2017) that has been established as a powerful and biologically plausible model to predict brain activity patterns in occipital cortex and the VVS (Doerig et al. 2023). A LMM model predicting rating response times by fixed effects of low-level (convolutional layers) and high-level (fully-connected layers) perceptual feature similarity yielded non-significant effects (convolutional: *Z* = −1.088, *χ*^2^ = 1.18, *P* = 0.277; fully-connected: *Z* = −0.967, *χ*^2^ = 0.94, *P* = 0.333), suggesting that response times for conceptual distance ratings did not depend on stimulus-dependent perceptual formats.

Similarly, we ensured that the effect of rated Euclidean distances on item memory confidence was not driven by the visual similarity of both images. A model predicting memory confidence by main effects of rated distance as well as DNN-derived distances between both images of a trial yielded a significant main effect of rated Euclidean distance (*Z* = 3.452, *χ*^2^ = 11.91, *P* < 0.001) but not low-level or high-level visual features (convolutional layers; *Z* = −0.626, *χ*^2^ = 0.39, *P* = 0.822; fully-connected layers; *Z* = 0.493, *χ*^2^ = 0.24, *P* = 0.886) suggesting that task-relevant but not stimulus-dependent visual features were relevant for subsequent item memory strength.

#### Exemplar and Concept discrimination performance

We repeated calculation of d-prime across confidence intervals split by new image categories. Specifically, we quantified exemplar discrimination (without new concepts) and concept discrimination (without new exemplars) which were both reliably above zero (exemplar discrimination: *M* = 0.625, *SD* = 0.226; *T*_45_ = 18.737, *P* < 0.001 | concept discrimination: *M* = 2.458, *SD* = 1.242; *T*_45_ = 13.425, *P* < 0.001). Comparison of exemplar and concept discrimination revealed better concept detection as compared to exemplar detection (AUC exemplar vs. concept: *T*_45_ = −21.414, *P* < 0.001) suggesting superior concept discrimination.

#### BOLD magnitude of Hits scales with behavioral confidence ratings

Extending univariate SME contrasts, we conducted an analysis testing whether the magnitude of BOLD activity for subsequently remembered images (Hits) scaled with behavioral confidence ratings. We performed this analysis separately for regions of the MTL and on the whole-brain level, by including a parametric regressor into first level models that weighed the magnitude of BOLD response for Hits by mean-centered memory confidence ratings. In the MTL, we found that HC (*T*_45_ = 2.949, *P* = 0.005), PRC (*T*_45_ = 4.371, *P* < 0.001) and PHC (*T*_45_ = 6.042, *P* < 0.001) but not EC (*T*_45_ = 0.695, *P* = 0.491) BOLD activity scaled with subsequent memory confidence ratings (fig. S6, left). On the whole-brain level, we found largely overlapping clusters with regions showing a positive SME as indicated by our previous analysis (fig. S6, right; Table S2).

#### Distance metrics within and across task-relevant spaces

We conducted follow up analyses to quantify the relationship between the behavioral distance metrics of concept relationships. Specifically, in a LMM where we modeled participants with random intercepts, a concept’s distance to the origin in task-relevant spaces was significantly associated with both rated trial-level distance (*Z* = 16.627, *χ*^2^ = 272.31, *P* < 0.001) and the z-scored number of neighbors after neighborhood density estimation (*Z* = −120.598, *χ*^2^ = 8286.05, *P* < 0.001). These relationships were confirmed by significant correlations across trials and participants between distance to origin and trial-level distance (*ρ* = 0.363, *P* < 0.001) as well as distance to origin and neighborhood density (*ρ* = −0.800, *P* < 0.001).

### Supplementary Figures

**Supplementary Figure 1:**
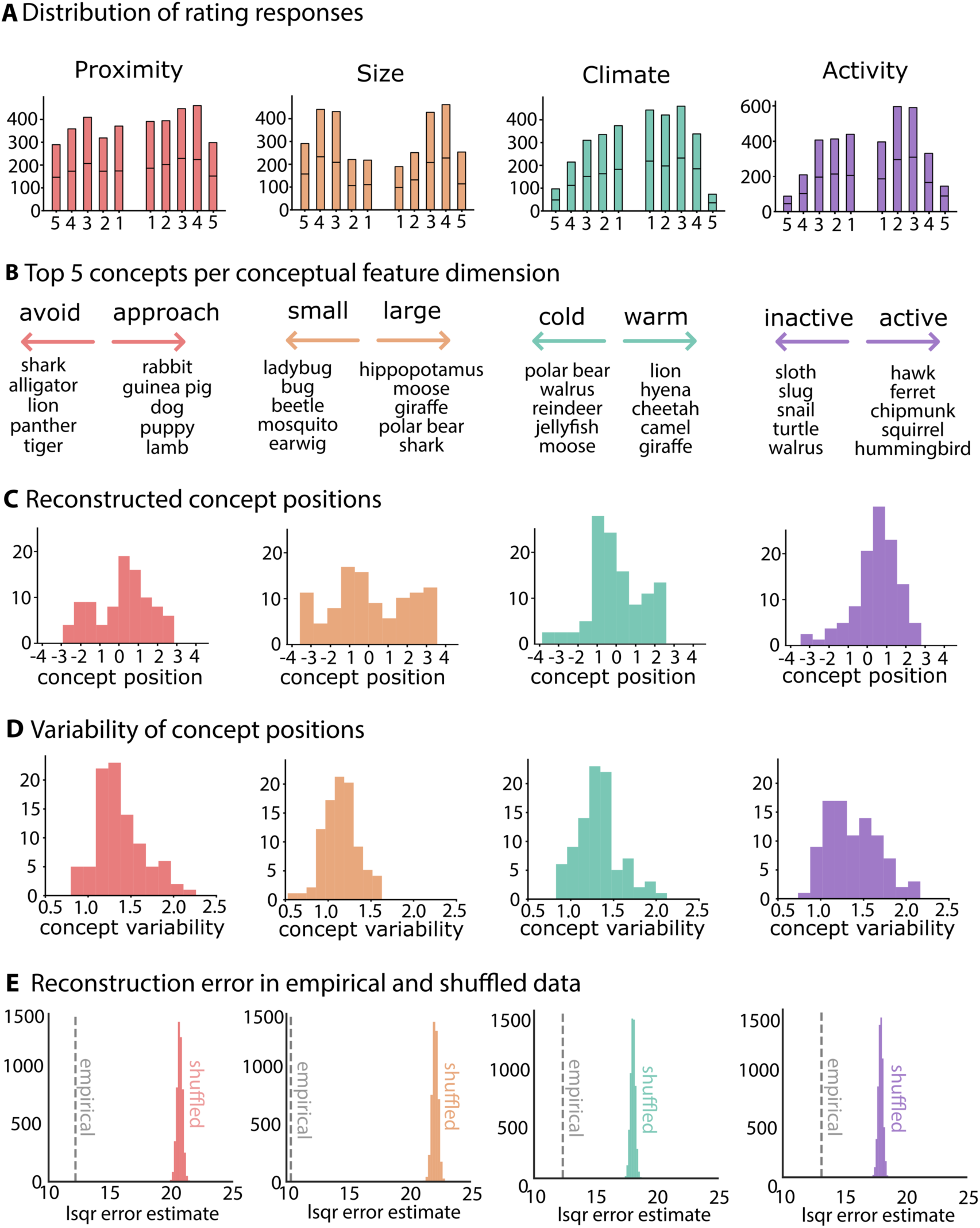
Rating responses and reconstructed concept positions across conceptual features. A) Histograms of rating responses (left hand, right hand) for each feature dimension (*Proximity*, *Size*, *Climate*, *Activity*). B) Top 5 concepts after dimension-specific reconstruction of concept positions across the entire sample of participants. C) Histograms of estimated concept positions after global space reconstructions. D) Histograms of the across-participant variability (i.e. standard deviation) of reconstructed concept positions. E) Validation of least-squares solutions by comparing the error of the empirical data (mean across participants) with the error of solutions obtained after shuffling empirical ratings within participants 5,000 times. Dashed vertical lines indicate the error of the empirical data. Colored histograms are distribution of error estimates after 5,000 permutation solutions, averaged across participants.

**Supplementary Figure 2:**
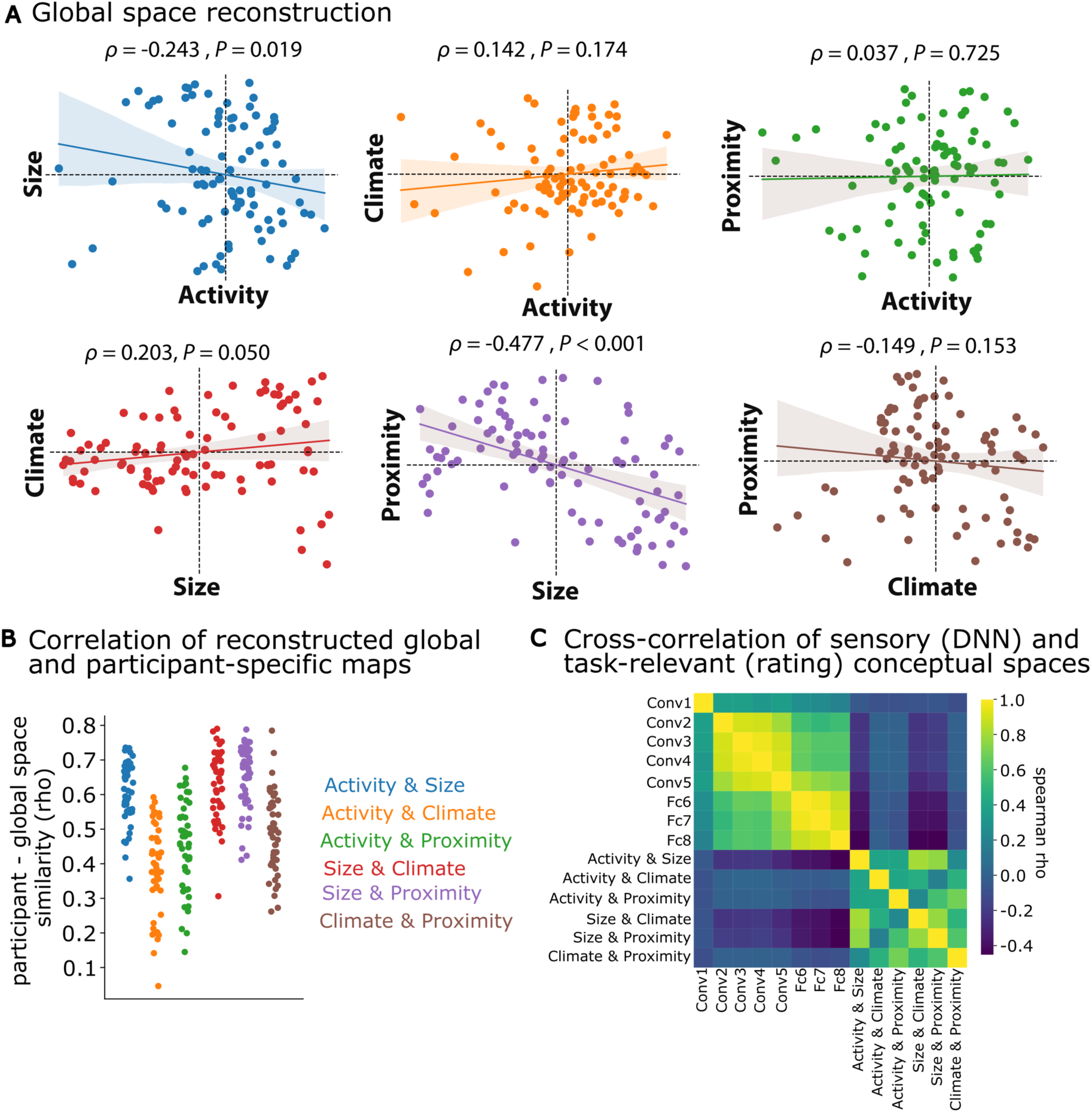
Global reconstruction of conceptual spaces. A) Reconstruction of representational spaces based on all ratings obtained in the experiment (i.e. global). Concept positions (dots) for each conceptual feature were grouped into all possible combinations of 2D spaces: *Size & Activity* (blue), *Activity & Climate* (orange), *Activity & Proximity* (green), *Size & Climate* (red), *Size & Proximity* (purple) and *Climate & Proximity* (brown). In each space, we correlated between concept positions of both feature dimensions using Spearman’s rank correlation. *P* values uncorrected. B) Cross-correlation matrix of representational geometries from DNN features (5 Convolutional layers: Conv1-Conv5; 3 Fully-connected layers: Fc6-Fc8) and global space reconstructions.

**Supplementary Figure 3:**
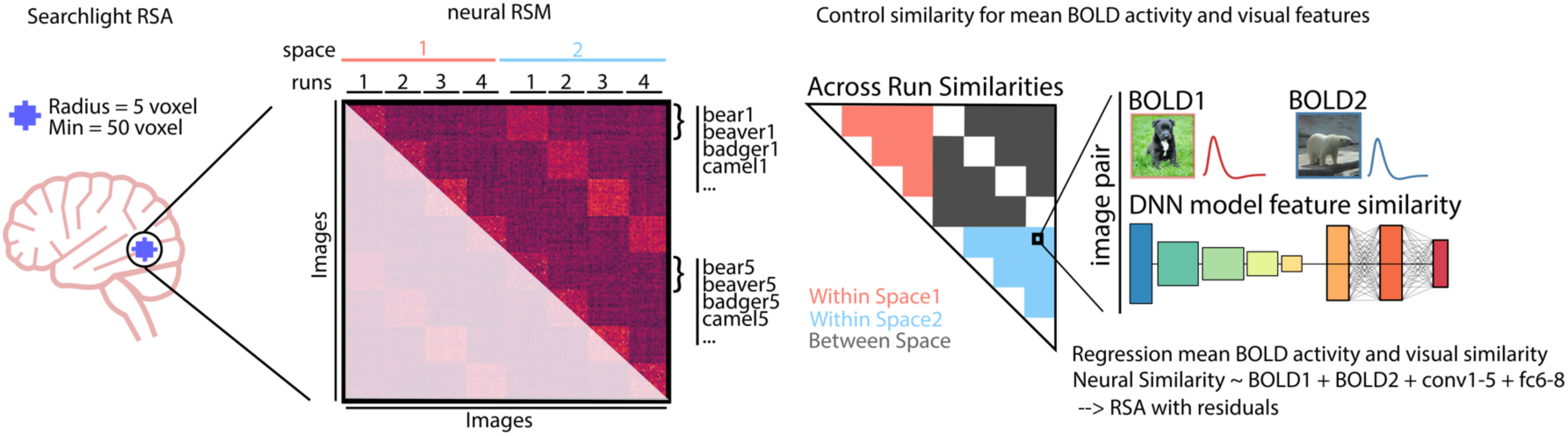
Searchlight RSA. Visual description of RSA / Searchlight analysis procedure. Across all voxels in a searchlight sphere / ROI we computed similarities between all image pairs, sorted the resulting symmetric image x image matrix according to space and run and performed all further steps on the upper triangle of the matrix (left). Then, we used between run similarities only to avoid temporal autocorrelations for images presented in the same run (middle). Finally, we regressed variance attributable to absolute levels of BOLD magnitude and perceptual similarity as predicted by DNN model layers from the neural similarity estimates (right).

**Supplementary Figure 4:**
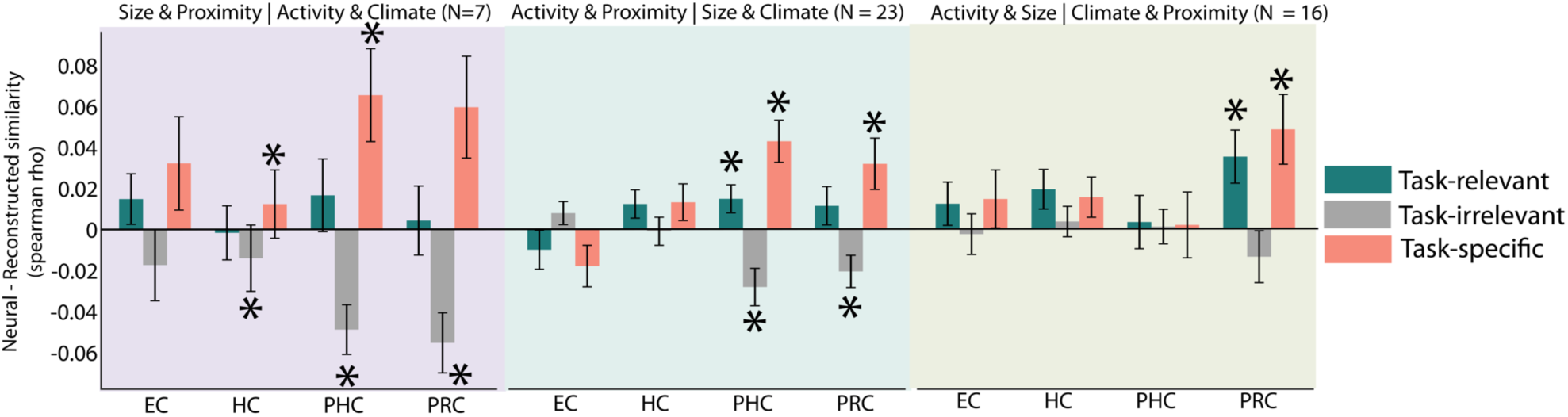
Task-dependent representational alignment of neural and reconstructed concept relationships per space. *N*=46 participants eligible for analysis grouped by feature dimensions that were explicitly rated. For each subgroup, average similarity of neural and reconstructed concept relationships for each subregion of the MTL are displayed, akin to analysis and results presented in Fig. 3A-C. * indicates *P* < 0.05 of one-sample *T* test, uncorrected.

**Supplementary Figure 5:**
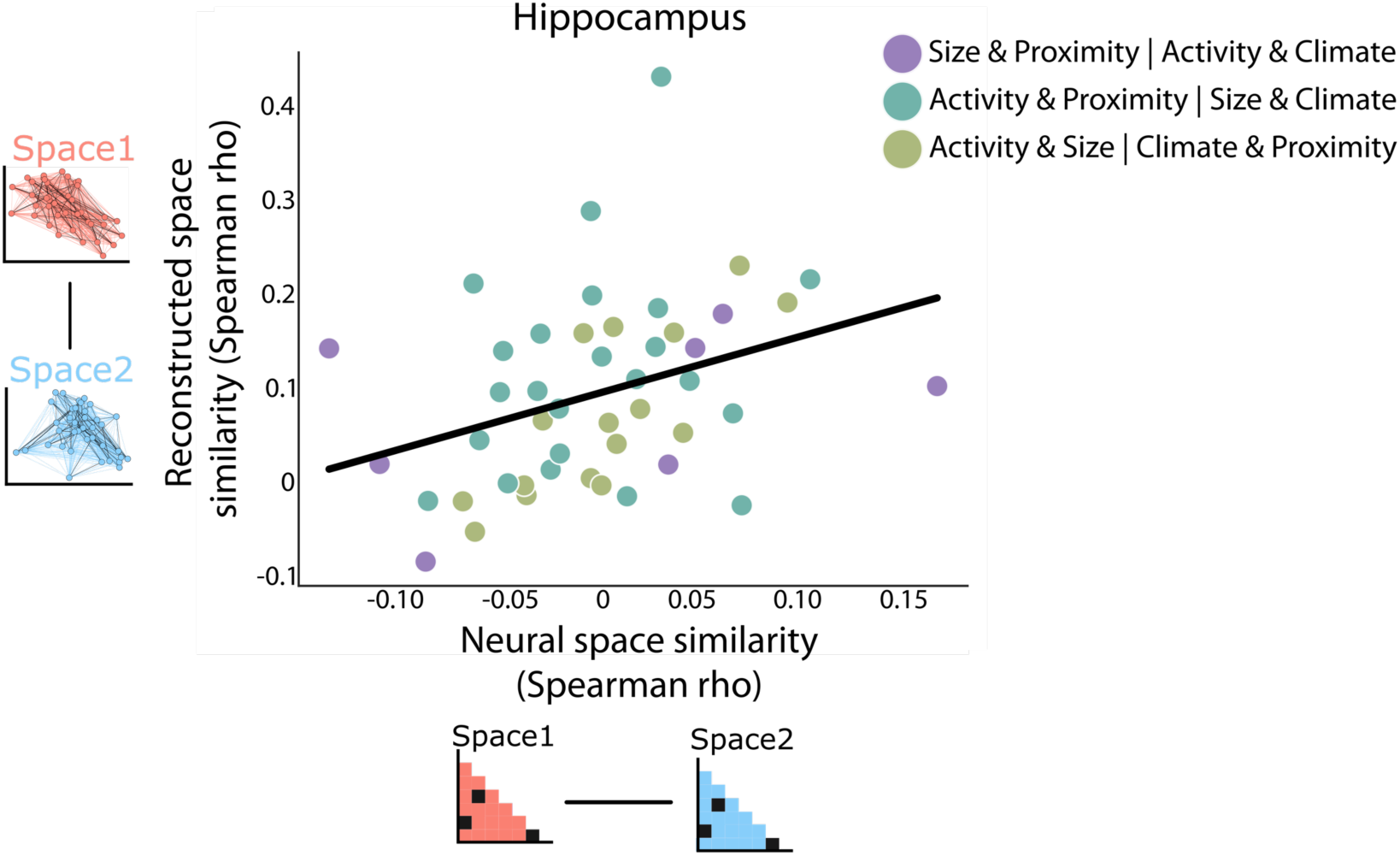
Space-level reorganization for subgroups of participants per rated spaces. Correlation of space1-space2 similarity across participants for neural (x-axis) and reconstructed (y-axis) concept relationships. Each dot is one participant, colored by which spaces were rated during the experiment (Purple: *Size* & *Proximity* | *Activity* & *Climate*, *N*=7, *ρ* = 0.357, *P* = 0.432; Olive: *Activity* & *Size* | *Climate* & *Proximity* - N=16, *ρ* = 0.783, *P* < 0.001; Teal: *Activity* & *Proximity* | *Size* & *Climate*, *N*=23, *ρ* = 0.208, *P* = 0.342).

**Supplementary Figure 6:**
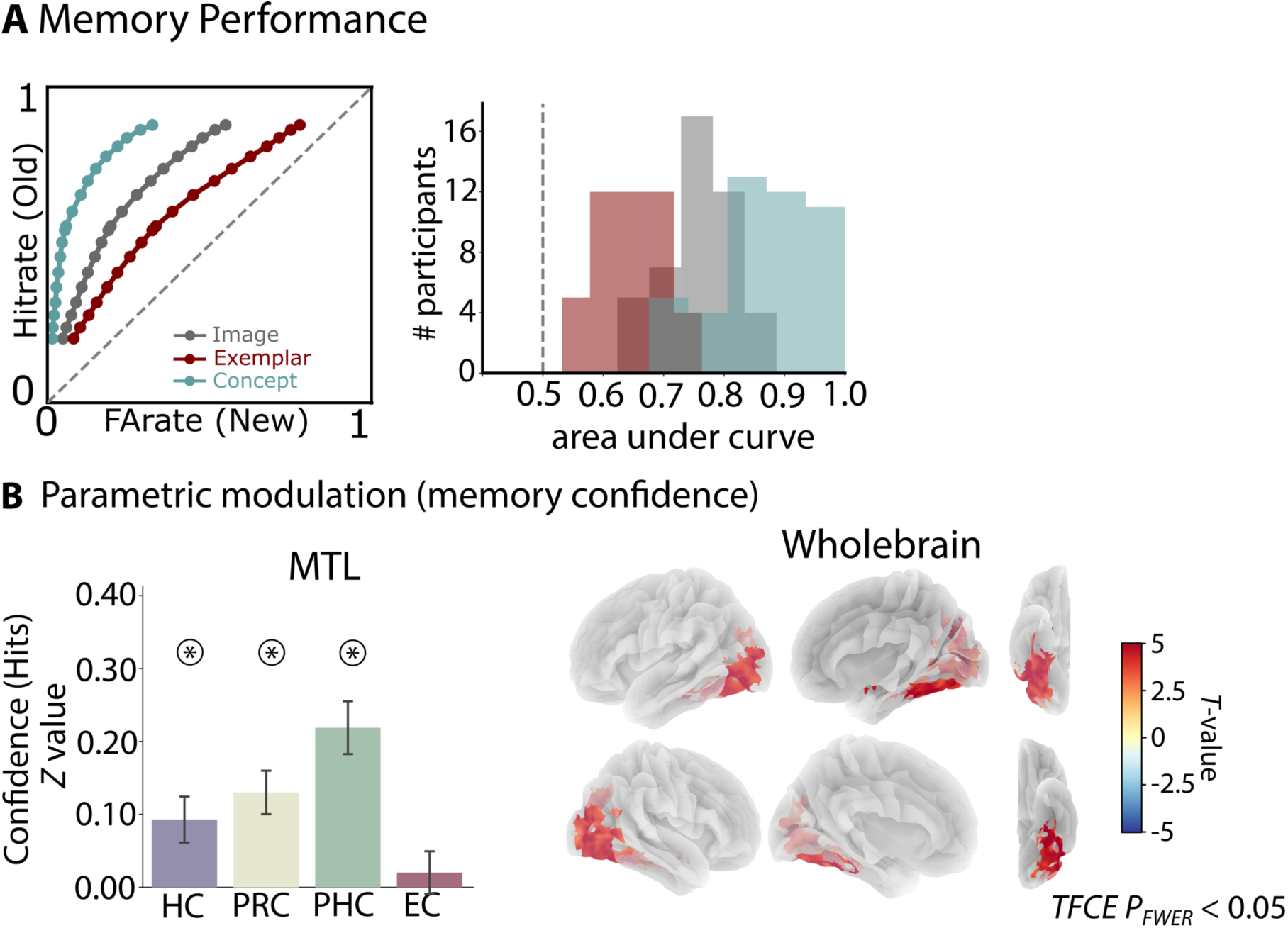
Exemplar vs. concept discrimination and parametric modulation of BOLD activity for hits. A) Left: Receiver-Operator Characteristics curves showing the ratio of hits vs. false alarms across the confidence scale (bins of 5 at scale from −50 to +50), separately for exemplar discrimination (without new concepts; red), concept discrimination (without new exemplars; teal) or both (grey). Right: histogram or area under the curve across participants. Dashed vertical line indicates chance level. B) Parametric modulation analysis, weighting BOLD activity of subsequently remembered images by behavioral confidence ratings for regions of the MTL (left) or on the whole-brain level (right). Whole-brain results thresholded at the cluster level after *TFCE* with *P*_FWER_ < 0.05.

**Supplementary Figure 7:**
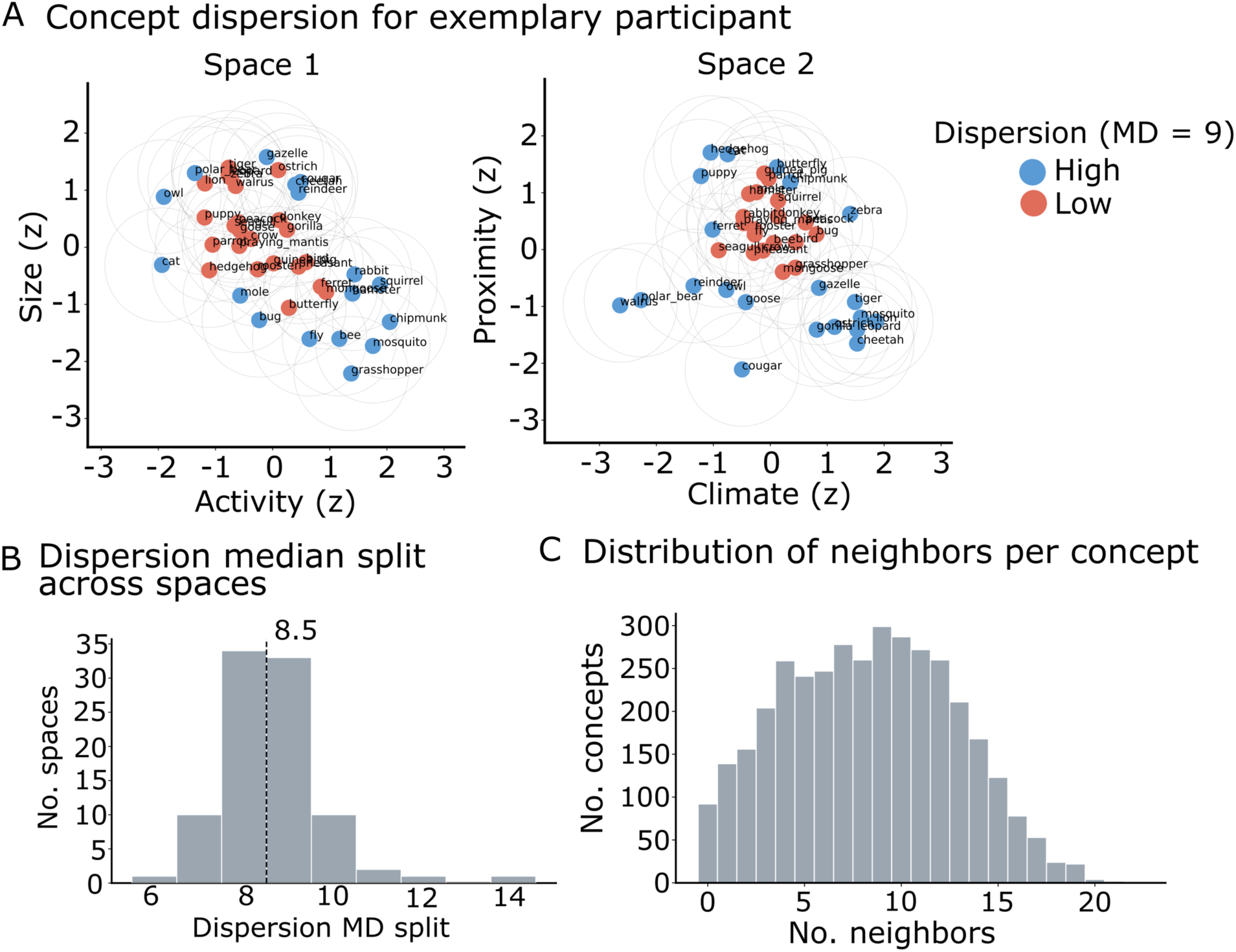
Concept dispersion. A) Reconstructed representational spaces of one example participant, with z-scored axes. Concepts are grouped into high (blue) and low (red) dispersion classes, indicating above or below median number of neighbors (*MD* = 9). Circles around each concept indicate search radius within which neighboring concepts are counted. B) Number of neighbors used for dispersion median split across spaces. C) Number of neighbors for all concepts across spaces.

**Supplementary Figure 8:**
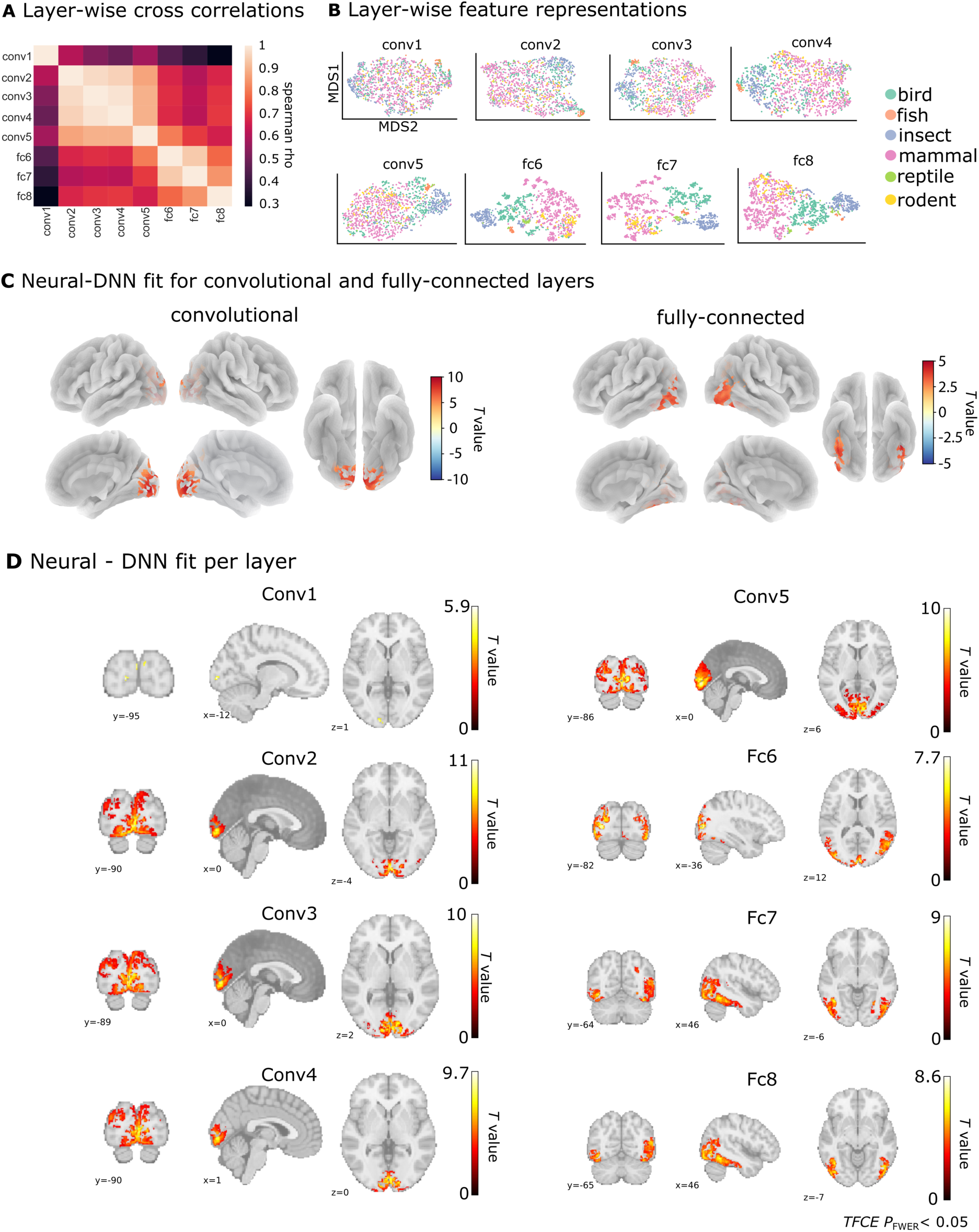
Layer-wise neural-DNN model fit. A) Cross-correlation matrix of pairwise layer similarities reflects similarity structure of the network. B) Two-dimensional *t-SNE* solutions for layer-wise feature similarities. Each dot represents a single image shown during the experiment. Dots are color coded by manual classification of animals into higher-level semantic animal categories: birds (green; e.g. cardinal, hawk), fish (orange; XY), insect (blue; bug, beetle), mammal (rose; e.g. monkey, gopher), reptile (green; praying mantis), rodent (yellow, e.g. mouse, squirrel). Feature-specific organization with increasing layer number reveals continuous transformation from low-level perceptual to semantic feature organization. C) Whole-brain neural-DNN model fit, averaged across convolutional layers (conv1-5; left) and fully-connected layers (fc6-8; right). D) Whole-brain searchlight results for the matching of neural and DNN layer representations across the brain. All clusters *P*_FWER_ < 0.05 after non-parametric permutation testing using *TFCE*.

**Supplementary Figure 9:**
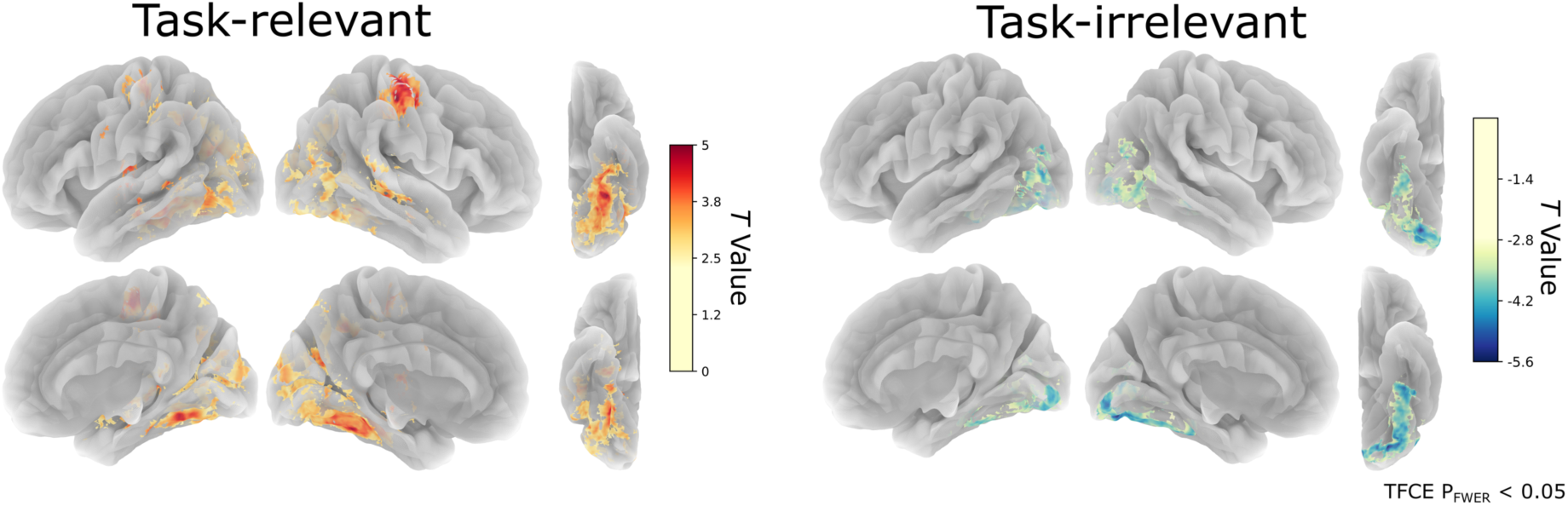
Task-related alignment between neural and reconstructed spaces. Wholebrain searchlight RSA results for testing the significance of neural and reconstructed concept relationships between task-relevant spaces (left; averaged similarities of space1-space1 | space2-space2) and task-irrelevant spaces (right: averaged similarities of space1-space2 | space2-space1). All colored voxels survive cluster correction after *TFCE* at *P*_FWER_ < 0.05. For cluster statistics see Table S5.

**Supplementary Figure 10:**
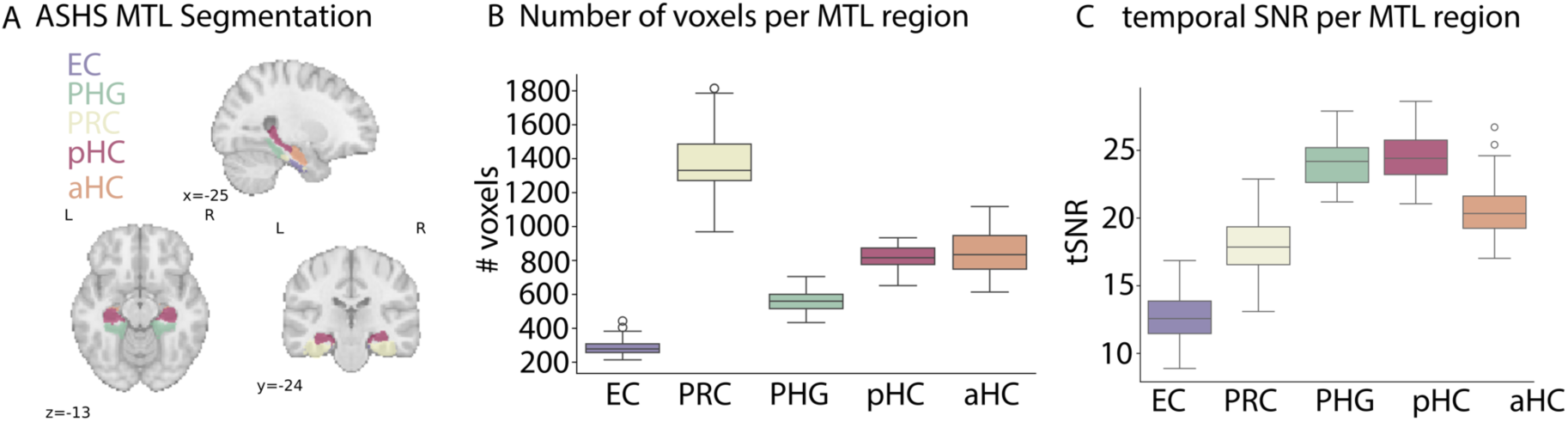
ASHS segmentation of medial temporal lobe. A) Segmentation of medial temporal lobe based on the ASHS atlas (ASHS-PMC-T1): entorhinal cortex (EC), parahippocampal gyrus (PHG), perirhinal cortex (PRC), posterior hippocampus (pHC) and anterior hippocampus (aHC). B) Mean number of voxels per ROI. C) Temporal signal-to-noise ratio (SNR) per region with 6mm smoothing. Error bars indicate standard deviation across participants.

### Supplementary Tables

**Supplementary Table 1:** Task-specific representational spaces in the MTL. *Searchlight-based RSA within the MTL of neural and rated representational spaces*

| Region | Contrast | ID | Location (ASHS) | Size (mm<sup>3</sup>) | x | y | z | T-Value | TFCE-Value |
| --- | --- | --- | --- | --- | --- | --- | --- | --- | --- |
| MTL | Task-relevant | 1 | PHC | 3576 | -31.0 | -36.6 | -16.1 | 5.10 | 9849 |
|  |  | 2 | HC (anterior) | 923 | 24.9 | -14.2 | -24.1 | 4.82 | 7071 |
|  | Task-irrelevant | 1 | PRC | 224 | 34.5 | -27.0 | -22.5 | -4.62 | -5769 |
|  |  | 2 | PRC | 89 | 34.5 | -17.4 | -30.5 | -4.34 | -5125 |
|  | Task-relevant vs. | 1 | PRC | 3591 | -31.0 | -28.6 | -19.3 | 5.55 | 11423 |
|  | Task-irrelevant | 2 | HC (posterior) | 772 | 29.7 | -25.4 | -14.5 | 4.97 | 6677 |
|  |  | 3 | PRC | 454 | 26.5 | -27.0 | -24.1 | 4.53 | 5899 |
|  |  | 4 | PRC | 217 | 34.5 | -17.4 | -27.3 | 4.33 | 5341 |

*Note.* All reported clusters with TFCE-based $P_{\text{FWER}} < 0.05$

**Supplementary Table 2:**
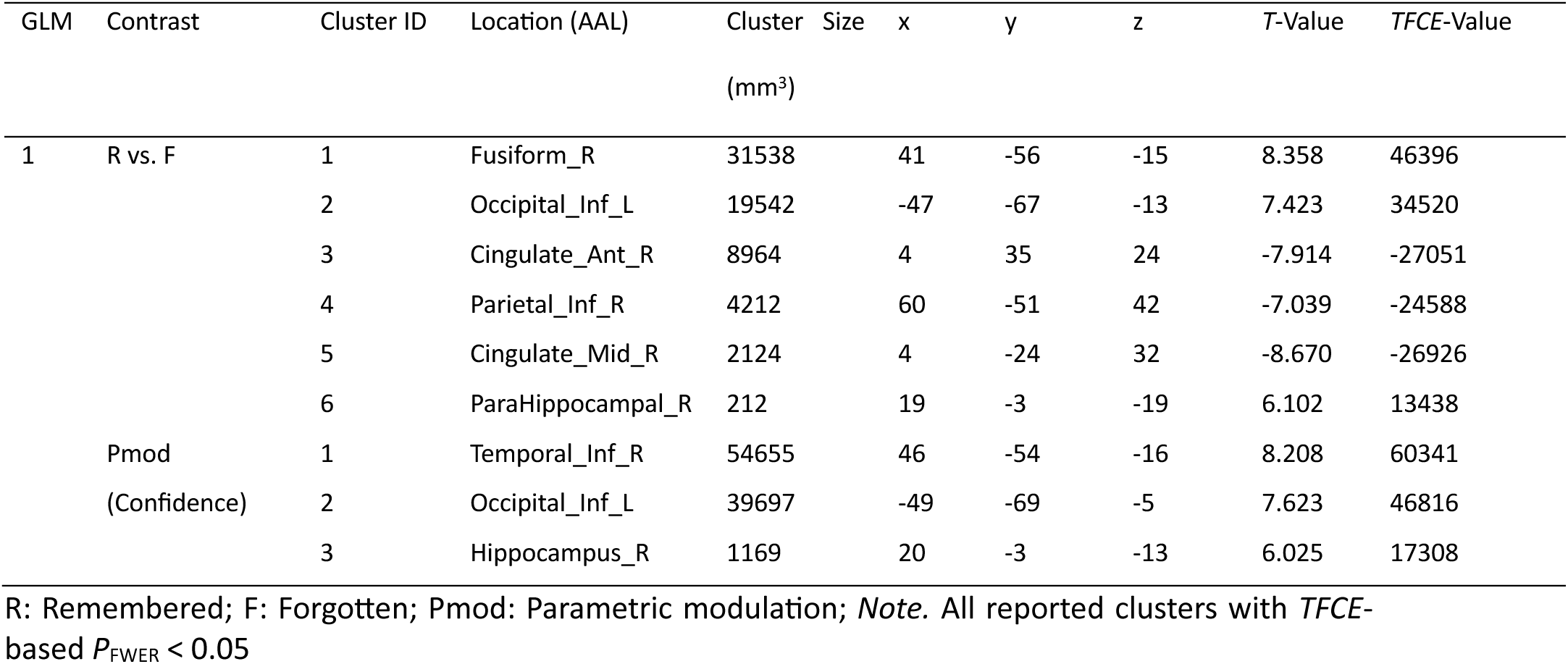
Cluster-statistics for univariate contrasts. *Univariate SME (remembered vs. forgotten) contrasts and parametric modulation (memory confidence)*

**Supplementary Table 3:**
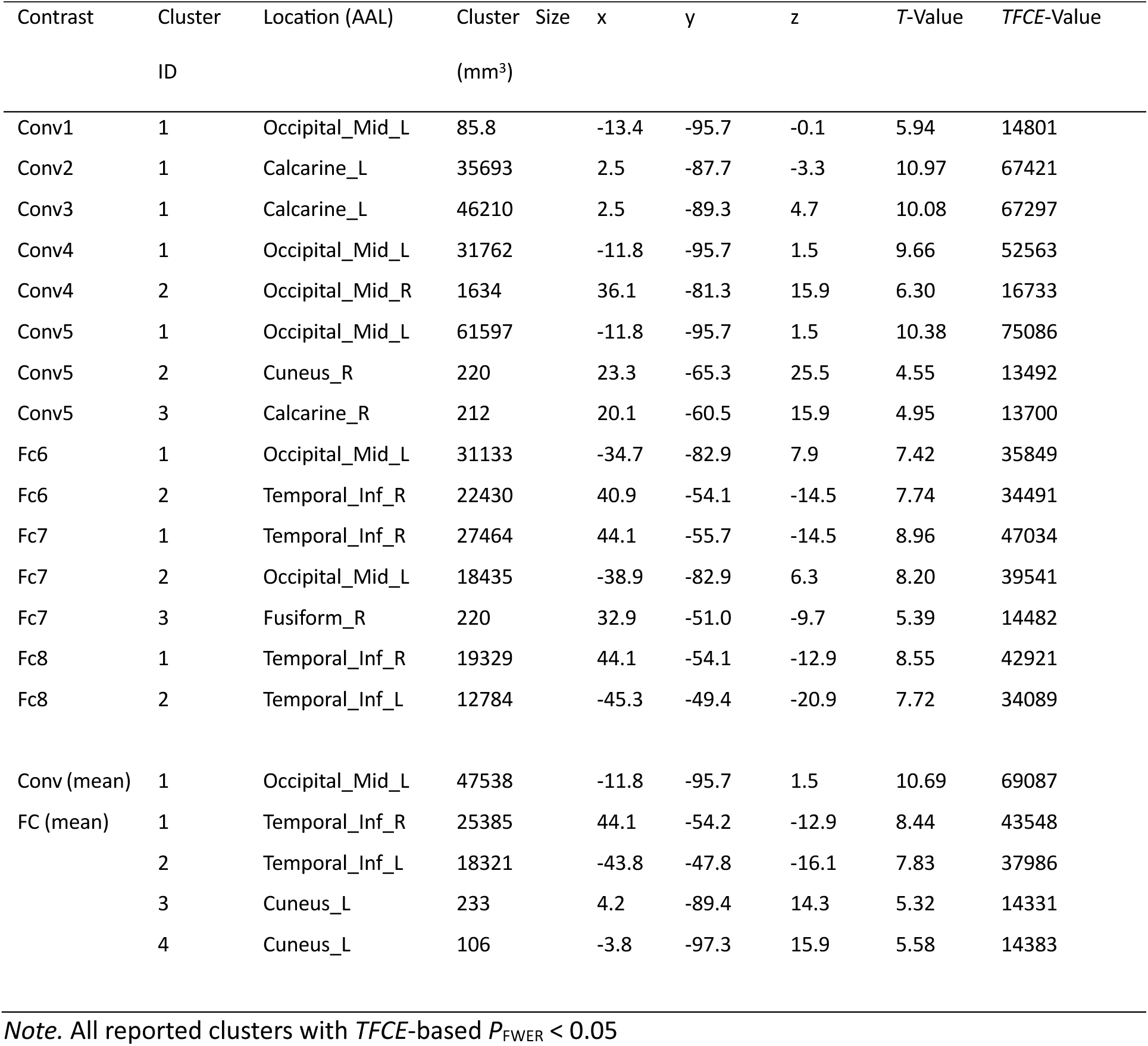
Layer-wise Neural-DNN fit. *Neural-DNN similarity, separately for each layer of the network as well as averaged layer-specific maps across convolutional (Conv) and fully-connected (Fc) layers*.

**Supplementary Table 4:**
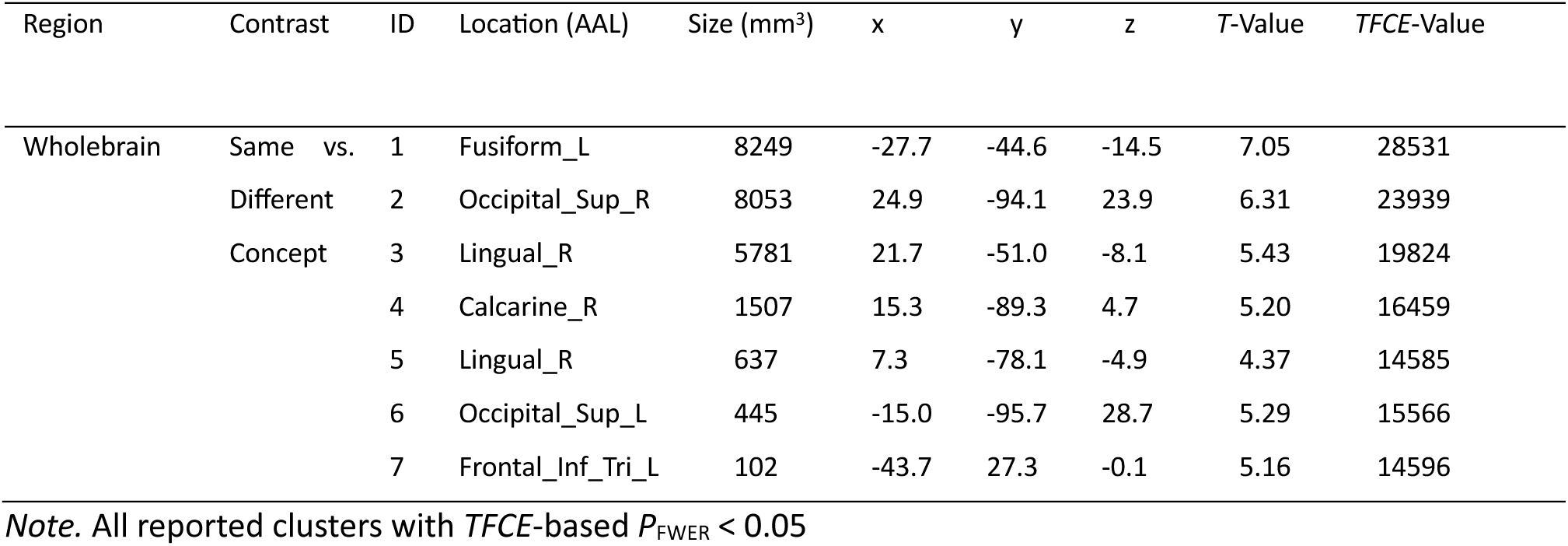
Concept-specific representational formats. *Conceptual formats RSA testing neural similarity across exemplars of the same vs. different animal*.

**Supplementary Table 5:** Task-specific representational spaces across the human brain. *Wholebrain Searchlight-based RSA of neural and rated representational spaces*

| Region | Contrast | ID | Location (AAL) | Size<br>(mm<sup>3</sup>) | x | y | z | T-Value | TFCE-Value |
| --- | --- | --- | --- | --- | --- | --- | --- | --- | --- |
| Wholebrain | Task-relevant | 1 | Fusiform_L | 100453 | -32.6 | -47.8 | 17.7 | 5.60 | 36368 |
|  |  | 2 | Postcentral_R | 10308 | 37.7 | -30.2 | 44.7 | 5.58 | 30513 |
|  |  | 3 | Temporal_Sup_R | 9442 | 58.5 | -31.8 | 1.5 | 5.31 | 24113 |
|  |  | 4 | ParaHippocampal_R | 4789 | 24.9 | -14.2 | -24.1 | 4.73 | 21980 |
|  |  | 5 | Precuneus_L | 3955 | -10.2 | -65.4 | 54.3 | 4.08 | 19036 |
|  |  | 6 | Precentral_L | 3738 | -42.2 | -23.8 | 62.3 | 5.24 | 22842 |
|  |  | 7 | Temporal_Sup_L | 2860 | -47.0 | -15.8 | -0.1 | 4.87 | 20892 |
|  |  | 8 | Temporal_Mid_L | 1553 | -49.0 | -39.8 | 1.5 | 4.61 | 19336 |
|  |  | 9 | Postcentral_L | 364 | -61.3 | -3.1 | 33.5 | 4.52 | 18259 |
|  |  | 10 | Angular_R | 110 | 44.1 | -70.2 | 31.9 | 3.66 | 17744 |
|  |  | 11 | Angular_R | 110 | 34.5 | -59.0 | 39.9 | 4.89 | 18038 |
|  | Task-irrelevant | 1 | Lingual_R | 44159 | 20.1 | -81.4 | -14.5 | -6.95 | -33444 |
|  |  | 2 | Occipital_Sup_R | 1197 | 26.5 | -95.7 | 20.7 | -3.97 | -16812 |
|  |  | 3 | Occipital_Mid_R | 183 | 39.3 | -86.2 | 9.5 | -3.93 | -16393 |
|  | Task-relevant vs.<br>Task-irrelevant | 1 | Fusiform_L | 215684 | -31.0 | -46.2 | -19.3 | 5.84 | 54998 |
|  |  | 2 | Postcentral_R | 17005 | 39.3 | -27.0 | 60.7 | 5.45 | 26774 |

*Note.* All reported clusters with *TFCE*-based $P_{\text{FWER}} < 0.05$

